# Impact of chromatin context on Cas9-induced DNA double-strand break repair pathway balance

**DOI:** 10.1101/2020.05.05.078436

**Authors:** Ruben Schep, Eva K. Brinkman, Christ Leemans, Xabier Vergara, Ben Morris, Tom van Schaik, Stefano G. Manzo, Daniel Peric-Hupkes, Jeroen van den Berg, Roderick L. Beijersbergen, René H. Medema, Bas van Steensel

**Affiliations:** Oncode Institute, Netherlands Cancer Institute, Amsterdam, the Netherlands; Division of Gene Regulation, Netherlands Cancer Institute, Amsterdam, the Netherlands; Division of Cell Biology, Netherlands Cancer Institute, Amsterdam, the Netherlands; Division of Molecular Carcinogenesis, Netherlands Cancer Institute, Amsterdam, the Netherlands; Robotics Screening Center, Netherlands Cancer Institute, Amsterdam, the Netherlands

## Abstract

DNA double-strand break (DSB) repair is mediated by multiple pathways, including classical non-homologous end-joining pathway (NHEJ) and several homology-driven repair pathways. This is particularly important for Cas9-mediated genome editing, where the outcome critically depends on the pathway that repairs the break. It is thought that the local chromatin context affects the pathway choice, but the underlying principles are poorly understood. Using a newly developed multiplexed reporter assay in combination with Cas9 cutting, we systematically measured the relative activities of three DSB repair pathways as function of chromatin context in >1,000 genomic locations. This revealed that NHEJ is broadly biased towards euchromatin, while microhomology-mediated end-joining (MMEJ) is more efficient in specific heterochromatin contexts. In H3K27me3-marked heterochromatin, inhibition of the H3K27 methyltransferase EZH2 shifts the balance towards NHEJ. Single-strand templated repair (SSTR), often used for precise CRISPR editing, competes with MMEJ, and this competition is weakly associated with chromatin context. These results provide insight into the impact of chromatin on DSB repair pathway balance, and guidance for the design of Cas9-mediated genome editing experiments.

## INTRODUCTION

The repair of DNA double strand breaks (DSBs) is crucial for genetic stability. In addition, it is a key step in CRISPR/Cas9-mediated genome editing (Jasin and Haber, 2016; Yeh et al., 2019). Several pathways can repair DSBs, including classical non-homologous end-joining (NHEJ), homologous recombination (HR) and microhomology-mediated end-joining (MMEJ) (McVey and Lee, 2008; Iliakis et al., 2015; Chang et al., 2017; Scully et al., 2019; Yeh et al., 2019). NHEJ directly re-joins blunt-ended DSBs, while HR typically uses the intact sister chromatid in G2 phase as a template to mend the break. In contrast, MMEJ recombines short homologous sequences that are close to either end of the DSB, and consequently results in a small deletion. An additional variant is single-stranded template repair (SSTR), a type of homology-directed repair that requires a single-stranded oligodeoxynucleotide (ssODN) donor sequence (Lin et al., 2014; Richardson et al., 2016). SSTR is highly relevant because it is leveraged in CRISPR/Cas9-mediated genome editing to generate precisely designed small mutations, such as point mutations or small insertions or deletions (indels) (DeWitt et al., 2016; Okamoto et al., 2019; Riesenberg et al., 2019).

Which pathway repairs a particular DSB depends in part on the local DNA sequence (Allen et al., 2018; Shen et al., 2018; Chakrabarti et al., 2019; Chen et al., 2019) and on the stage of the cell cycle (reviewed in Chapman et al., 2012; Hustedt and Durocher, 2016; Mladenov et al., 2016). In addition, local chromatin packaging can affect the choice of repair pathway (Jeggo and Downs, 2014; Clouaire and Legube, 2015; Kalousi and Soutoglou, 2016; Scully et al., 2019). Most studies of chromatin effects so far have focused on the balance between HR and NHEJ. For example, the histone modification H3K36me3, which is present along active transcription units, is thought to promote HR (Daugaard et al., 2012; Aymard et al., 2014; Carvalho et al., 2014; Pfister et al., 2014; Clouaire et al., 2018). Paradoxically, H3K9 di- or trimethylated (H3K9me2/3) heterochromatin, which packages transcriptionally inactive regions of the genome, has also been implicated in promoting HR (Sun et al., 2009; Baldeyron et al., 2011; Lee et al., 2013; Soria and Almouzni, 2013; Alagoz et al., 2015), although some single-locus studies in mouse and fruit fly found no major change in the balance between NHEJ and HR when a sequence was shifted between heterochromatin and euchromatin states (Janssen et al., 2016; Kallimasioti-Pazi et al., 2018). Furthermore, reduced binding of HR proteins was observed at a locus that was artificially tethered to the nuclear lamina (Lemaitre et al., 2014), suggesting that spatial positioning of the DSB inside the nucleus may also play a role.

Much less is known about the impact of chromatin on MMEJ and SSTR. Like HR, these pathways require resection of the DNA ends to produce single-stranded DNA overhangs, but downstream of this step the mechanisms and responsible proteins diverge (Chang et al., 2017; Scully et al., 2019; Yeh et al., 2019). It is thus possible that the local chromatin environment also modulates MMEJ and SSTR in unique ways, but this has remained largely unexplored (Clouaire and Legube, 2019; Mitrentsi et al., 2020).

One strategy to investigate the impact of local chromatin context on repair pathway balance is to generate DSBs at various genomic locations with known chromatin states, and compare pathway utilization across these locations (van Overbeek et al., 2016; Clouaire et al., 2018; Chakrabarti et al., 2019). However, with such an approach it is difficult to separate the effects of chromatin context from the effects of sequence context, because both vary simultaneously along the genome. Ideally, different chromatin contexts are compared while the sequence context is kept fixed.

Here, we report a strategy that effectively tackles these challenges in human cells. The strategy consists of two parts. First, we developed a reporter that, when cut with Cas9, produces distinct “scars” when repaired by either NHEJ, MMEJ or SSTR; high-throughput sequencing of these scars provides highly accurate measurements of the relative activities of the three pathways. Second, we used a modification of our TRIP method (Akhtar et al., 2013) to insert this reporter into >1,000 random genomic locations, tracking each individual reporter in parallel by molecular barcoding. We thus systematically measured the relative activities of NHEJ, MMEJ and SSTR as function of chromatin context in >1,000 genomic locations.This yielded unique datasets that (1) comprehensively sample the broad diversity of chromatin ‘flavors’ across the entire genome; (2) bypass the confounding effects of varying sequence context; (3) probe three of the most relevant pathways, with high accuracy and sensitivity. The results provide a detailed view of the impact of chromatin context on the relative activities of the three repair pathways.

## RESULTS

### Multiplexed DSB repair pathway assay: principle

We developed a strategy to measure the relative activity of several DSB repair pathways in more than one thousand genomic locations that include all known common chromatin states. The strategy employs a pathway-specific reporter construct that contains a short DNA sequence (derived from the human *LBR* gene) that predominantly produces a +1 insertion or a −7 deletion when cut at a specific base pair (bp) position by Cas9 (**Figure 1A**). We previously found that these two indels are primarily the result of NHEJ (+1) and MMEJ (−7), respectively (Brinkman et al., 2018) and we provide additional support below. The relative abundance of these signature indels can therefore be interpreted as a measure of the relative activity of these two pathways. Furthermore, this readout can be extended to include SSTR (see below). We note that HR cannot be detected with this assay, because HR generally repairs DSBs perfectly, and perfectly repaired DNA cannot be distinguished from uncut DNA. However, we previously estimated that perfect repair of this reporter sequence is rare, and by inference that the contribution of HR is likely to be very minor (Brinkman et al., 2018).

**Figure 1:**
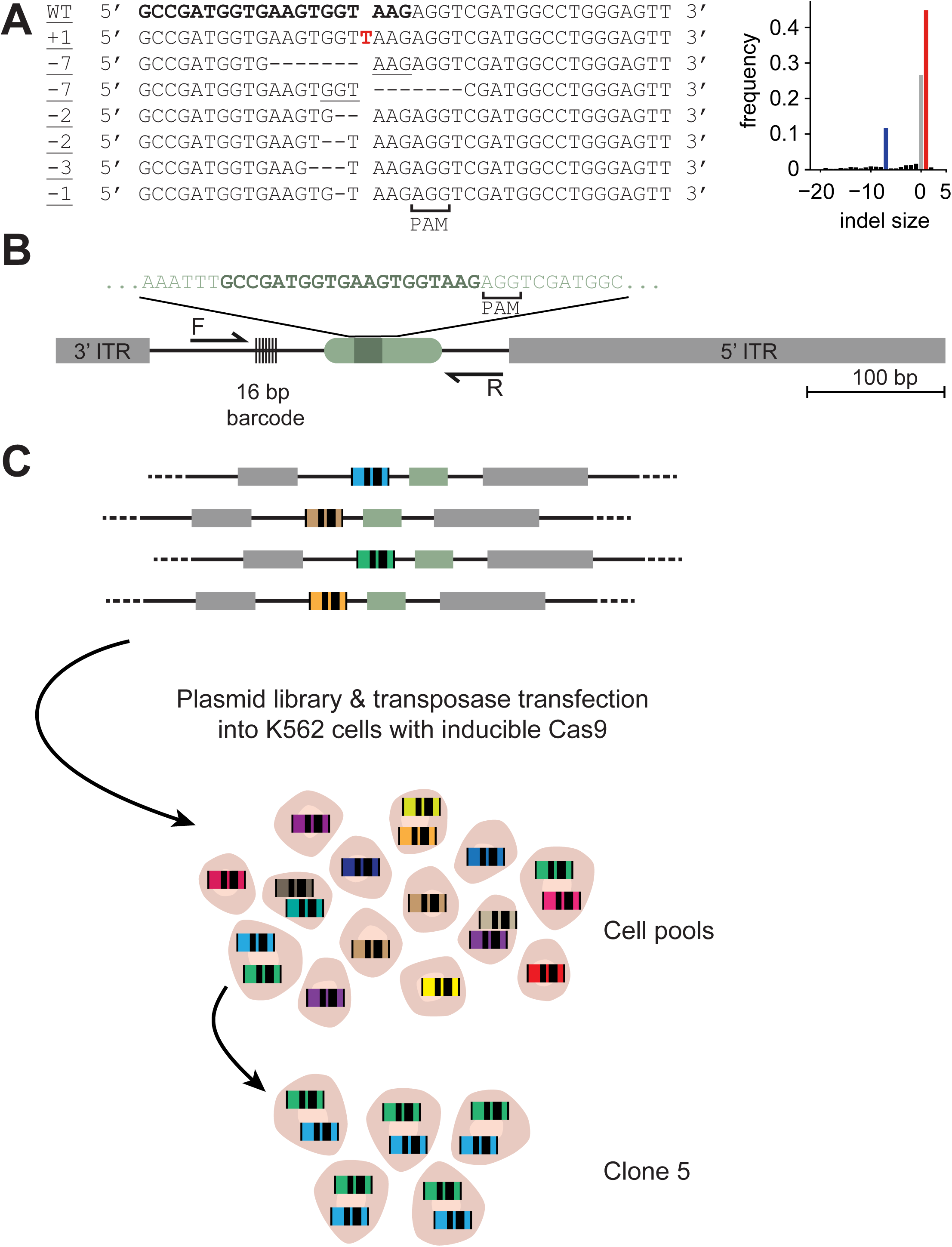
Principle of multiplexed DSB repair pathway reporter assay. **(A)** Left panel: sequences of the most common insertions and deletions produced after break and repair with induced with LBR2 sgRNA. Inserted nucleotide in red, microhomologies are underlined. Right panel: indel frequency distribution after cutting the *LBR* gene. Negative indel sizes refer to deletions, positive sizes refer to insertions. The +1 and −7 indels are marked in red and blue, respectively. **(B)** Schematic of the TRIP construct. ITR, inverted terminal repeat of PiggyBac transposable element; the LBR gene-derived sequence fragment is shown in light green, with the sgRNA target sequence in dark green. PCR primers are indicated by the arrows (F and R). **(C)** Schematic of the TRIP experimental setup. See main text.

With this reporter, we implemented a variant of the TRIP technology (Akhtar et al., 2013) to systematically probe the effects of many chromatin environments on the repair pathway usage. We inserted the reporter sequence into a PiggyBac transposon vector, together with a 16 bp random barcode sequence that was located 56 bp from the DSB site (**Figure 1B**). We then randomly integrated this construct into the genomes of pools of K562 cells (**Figure 1C**). We chose K562 cells because the chromatin landscape has been extensively characterized (**Table S1-2**) (Encode Project Consortium, 2012; Schmidl et al., 2015; Schwalb et al., 2016; Chen et al., 2018; Ott et al., 2018; Shah et al., 2018). From this pool, we also generated a number of clones for smaller scale experiments. Each copy of the integrated reporter carried a different barcode. We mapped the genomic locations of these Integrated Pathway Reporters (IPRs) together with their barcode sequences by inverse PCR (Akhtar et al., 2013). Next, after Cas9-mediated DSB induction and the ensuing repair, we determined the accumulated spectrum of indels of each individual IPR in a multiplexed fashion, by PCR amplification (see primer locations in **Figure 1C**) followed by high-throughput sequencing. Because each barcode is linked to its genomic location, the location specific sequence information enabled us to infer the relative DSB repair pathway usage at each location. Comparison of the resulting data to the local chromatin state of the IPRs then provides insight into the impact of chromatin context on DSB repair pathway usage.

### Implementation and validation of the multiplexed reporter assay

For these experiments we used a human K562 cell line that expresses Cas9 protein in an inducible manner (Brinkman et al., 2018). We generated two cell pools with 979 and 1099 (total 2078) uniquely mapped IPRs (**Figure 2A; Figure S1A**). In addition, we established 14 clonal cell lines, for which the barcodes and locations were also mapped. On average these clones carried 6.8 integrations, which we take as an estimate of the numbers of integrations per cell in the cell pools. For some additional analyses described below, we used one clone (clone 5) with 19 mapped IPRs that are located across most major chromatin types (**Figure S1A, B**).

**Figure 2:**
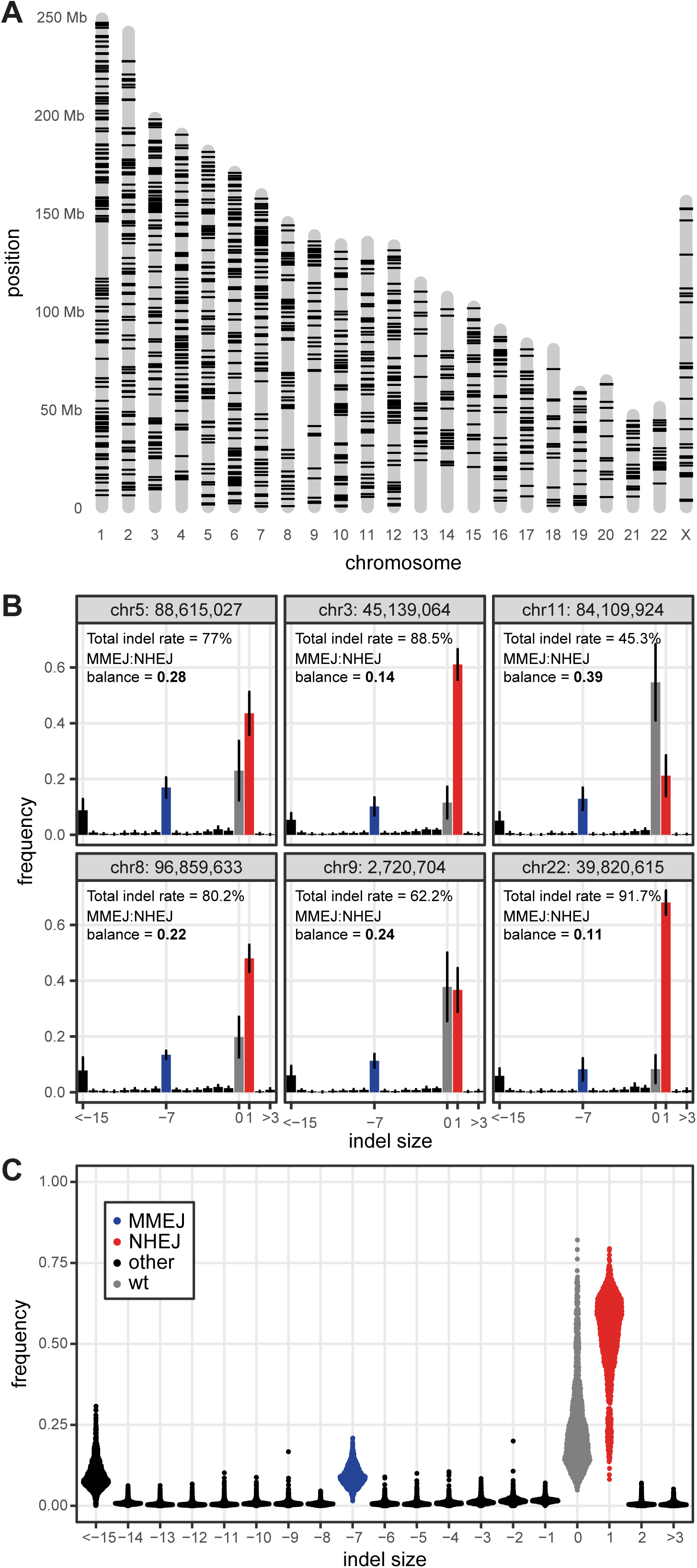
Multiplexed detection of DSB repair pathway usage. (**A**) Genomic integration coordinates of 1229 uniquely mapped IPRs (both cell pools combined) that passed filtering as described in the methods and are used in this work. (**B**) Indel frequency distributions of six randomly selected IPRs, 64 hours after Cas9 induction. Data are average of 6 independent replicates. Error bars are ± SD. Gray: wild-type sequence, red: +1 insertion, diagnostic of NHEJ, blue: = −7 deletion, diagnostic of MMEJ, black: other indels. (**C**) Indel frequencies of all IPRs shown in **A**, 64 hours after Cas9 induction. Data are average of 2-6 independent replicates.

Next, we induced Cas9 in the cell pools by ligand-dependent stabilization (Banaszynski et al., 2006) and transfection with the sgRNA (named sgRNA-LBR2). We collected genomic DNA after 64 hours and determined the indel spectra of all IPRs. At this time point, indel accumulation in the *LBR* gene has reached near-saturation (Brinkman et al., 2018). After applying stringent quality criteria (see Methods) we obtained robust indel spectra of 1229 IPRs (**Figure S1A**); the other IPRs were mostly discarded because they were insufficiently represented in the cell pools. Overall, the IPRs of both pools showed a similar pattern of indels as the endogenous *LBR* sequence in the same cells, dominated by +1 and −7 indels (**Figure 2B, C**; **Figure S1C, D**). This supports previous findings that the sequence determines the overall indel pattern (van Overbeek et al., 2016; Allen et al., 2018; Shen et al., 2018; Chakrabarti et al., 2019; Chen et al., 2019), but we also observe clear variations in indel frequencies.

As noted before (Brinkman et al., 2018), the −7 deletions come in two variants that both involve 3-nucleotide microhomologies (**Figure 1A**), consistent with MMEJ. To further verify that the +1 and −7 indels indeed represent NHEJ and MMEJ, respectively, we depleted or inhibited several pathway-specific proteins (Chang et al., 2017; Scully et al., 2019) in either the pools or clone 5 (**Figure S2**). The +1 insertion was strongly reduced and the −7 deletion was increased by inhibition of DNA-PKcs by the compounds NU7441 (**Figure S2A, B**) or M3814 (**Figure S2C**). DNA-PKcs is a key component of NHEJ (Gottlieb and Jackson, 1993). In contrast, the −7 deletions but not the +1 insertion were selectively reduced upon depletion of DNA polymerase theta (POLQ) and CtIP, which are proteins of the MMEJ pathway (Sartori et al., 2007; Chan et al., 2010; Mateos-Gomez et al., 2015) (**Figure S2D, E**). Knockdown of Rad51 also caused a slight reduction in −7 deletions with a negligible change in +1 insertions (**Figure S2D, E**), while in yeast Rad51 inhibits MMEJ activity by promoting HR instead (Villarreal et al., 2012; Deng et al., 2014). Aside from this latter experiment, all other evidence indicates that the +1 and −7 indels are primarily the result of NHEJ and MMEJ, respectively.

### Effects of chromatin context on overall indel frequencies

Using the indel spectra from the two cell pools, we first investigated the impact of chromatin context on total indel frequencies (TIF; i.e., the proportion of reporter sequences carrying any type of indel). Across the IPRs these frequencies varied from ∼25% to ∼100% (**Figure 3A, B**). This variation most likely reflects differences in either the cutting efficiency by Cas9, or the DSB repair rate, or both. We repeated these experiments with three different sgRNAs targeting different sequences in the same reporter (**Figure 3A, B**). This yielded overall indel frequencies that strongly correlated with sgRNA-LBR2 and with each other, although with sgRNA-LBR2 we may have approximated saturation of indels more than with the other sgRNAs (**Figure 3B, C**).

**Figure 3:**
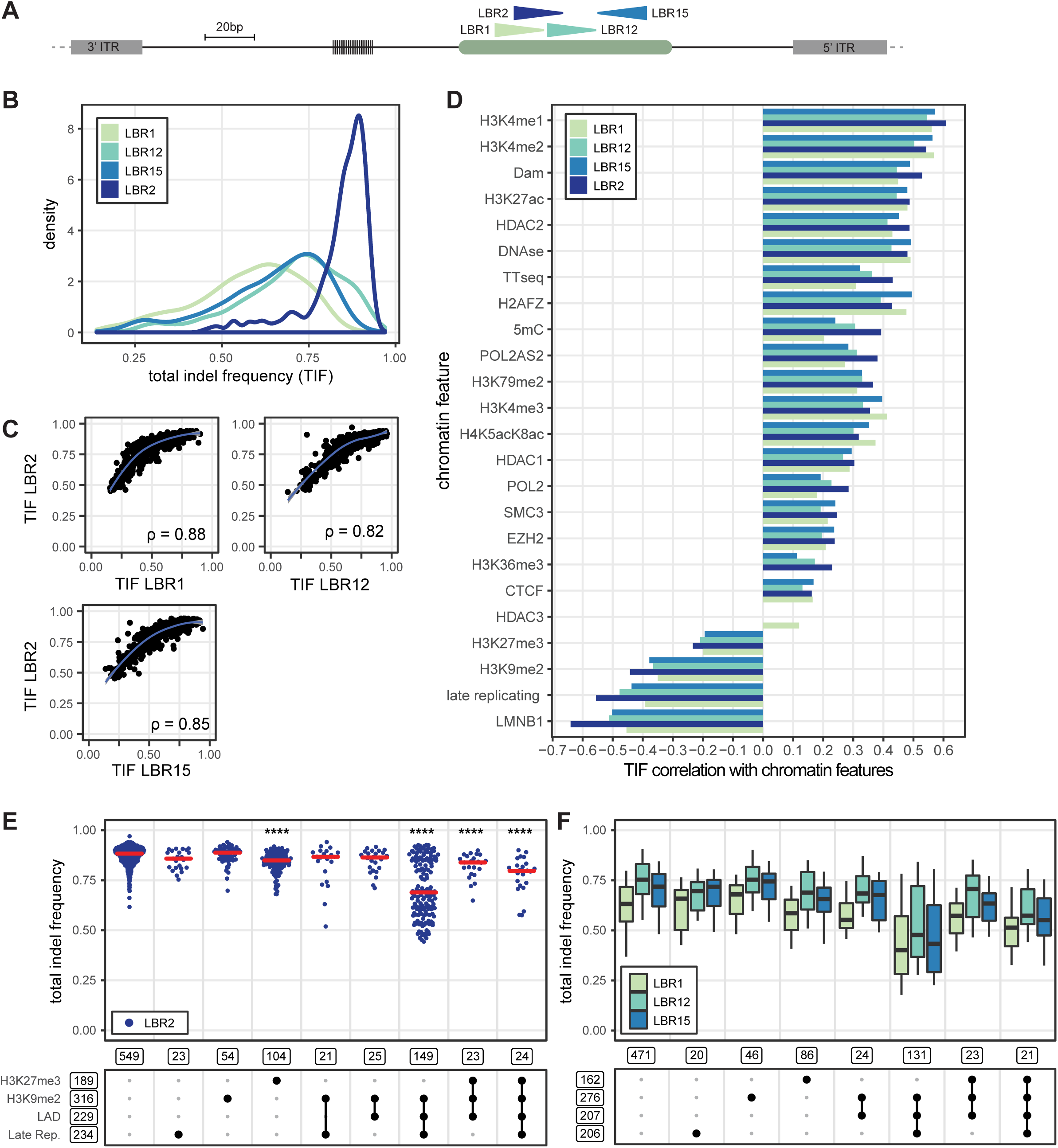
IPR total indel frequency varies as a function of chromatin context. (**A**) Schematic of the IPR with four different gRNA target sites indicated by the arrow heads. The arrow heads point toward their PAM site. (**B**) Total indel frequency distributions of IPRs after targeting by four different sgRNAs. For each sgRNA, IPRs were only included if they yielded reliable data in at least two independent experiments (LBR2: 1010 IPRs [n = 2-8]; LBR1: 956 IPRs [n = 2]; LBR12: 942 IPRs [n = 2]; LBR15: 932 IPRS [n = 2]). (**C**) Scatter plots of total indel frequencies obtained with LBR2 versus the three other sgRNAs. rho is Spearman’s rank correlation coefficient. (**D**) Pearson’s correlations between total indel frequencies in IPRs and the local intensities of 24 chromatin features at the IPR integration sites, for four different sgRNAs. Only statistically significant correlation values (p < 0.001) are shown. Chromatin features are ordered by the LBR2 correlation coefficients. (**E**) Total indel frequency at each IPR obtained with LBR2 sgRNA, split into different combinations of heterochromatin features present, as indicated by black dots in the scheme below the graph. Boxed numbers indicate the number of IPRs in each group; only groups with >20 IPRs are shown. Asterisks mark p-values according to the Wilcoxon test, compared to euchromatin IPRs (most left column): * p < 0.05, ** p < 0.01, *** p < 0.001, **** p < 0.0001. (**F**) Same as **E**, but boxplots for the other three sgRNAs. The boxes represent 75% confidence interval, the horizontal line within represents the median, the error bars represent and 95% confidence intervals.

Because the sequences of the IPRs are identical (except for the short barcodes located 56 bp from the cut site), the differences in indel frequencies across integration sites are presumably due to variation in the local chromatin environment. To investigate this, we correlated the indel data for each sgRNA with a curated set of 24 genome-wide maps of chromatin features that represent most of the known main chromatin types (**Table S1-2**). The IPRs lack gene regulatory elements and are only 640 bp long, and may thus be expected to adopt the local chromatin state. Chromatin immunoprecipitation experiments for three histone modifications generally confirmed this (**Figure S3**). This is consistent with previous studies showing that integrated reporters adopt and strongly respond to the local chromatin state (Akhtar et al., 2013; Corrales et al., 2017; Leemans et al., 2019). We therefore assume that the chromatin state of the integration positions is a reasonable approximation of the chromatin state of the IPRs themselves.

Overlaying of the indel frequencies with the 24 chromatin maps revealed that overall indel frequencies generally correlated positively with various markers of euchromatin and negatively with markers of heterochromatin. These correlations were highly consistent between the four sgRNAs that we tested (**Figure 3D**). On average, IPRs integrated in heterochromatin regions showed lower indel frequencies than in euchromatic regions. However, within heterochromatin the magnitude of this effect varied depending on the specific combination of features (**Figure 3E, F**). The most pronounced effect was observed in regions marked by the combination of H3K9me2, lamina-associated domains (LADs), and late replication. In these regions the distribution of indel frequencies appears to be bimodal, suggesting that another unknown feature plays an important role as well. Remarkably, when H3K27me3 is additionally present the reduction of indel frequencies is less pronounced, and regions marked by H3K27me3 alone only slightly affect indel frequencies compared to euchromatin. Thus, H3K27me3 only mildly impedes Cas9 editing and may even counteract the effects of other heterochromatin features. Regions marked by H3K9me2 together with either late replication or lamina interactions (but not both) show only marginally reduced indel frequencies, compared to the triple-marked regions (**Figure 3E, F**). For euchromatin regions, we did not survey combinatorial effects, because there are too many possible combinations and hence statistical power is insufficient.

Together, these results indicate that the overall indel frequency depends on the local chromatin context, and that heterochromatin features are correlated with the efficiency of indel accumulation in a combinatorial manner. These effects may be through modulation of Cas9 cutting efficiency, modulation of indel-forming repair rates, or both.

### Impact of chromatin context on MMEJ:NHEJ balance

Next, we analyzed the variation in the *balance* between MMEJ and NHEJ, throughout this paper referred to as “MMEJ:NHEJ balance” and defined for each IPR as the number of −7 reads over the sum of −7 and +1 reads. This balance varied profoundly depending on the integration site (**Figure 4A, B; S4A**). Strikingly, NHEJ activity correlated generally positively with markers of euchromatin. NHEJ showed the strongest positive correlations with H3K4me1, H3K4me2 and H3K27ac, which are histone modifications that primarily mark enhancers and to a lesser extent promoters (Gasperini et al., 2020). The strongest negative correlations of NHEJ included multiple markers of heterochromatin such as H3K27me3, H3K9me2 and LADs (**Figure 4C**; **S4B**). MMEJ showed the inverse relationships (**Figure 4C**; **S4B**). Within heterochromatin we further explored whether certain combinations of chromatin features are more predictive than others. The strongest effect on the balance was observed in IPRs located in regions marked by the combination of H3K9me2, late replication and LADs (**Figure 4D, S4C**). Regions marked by two of these three features showed less pronounced but significant increases in MMEJ:NHEJ balance compared to euchromatin regions, as did regions marked by H3K27me3 (**Figure 4D, S4C**).

**Figure 4:**
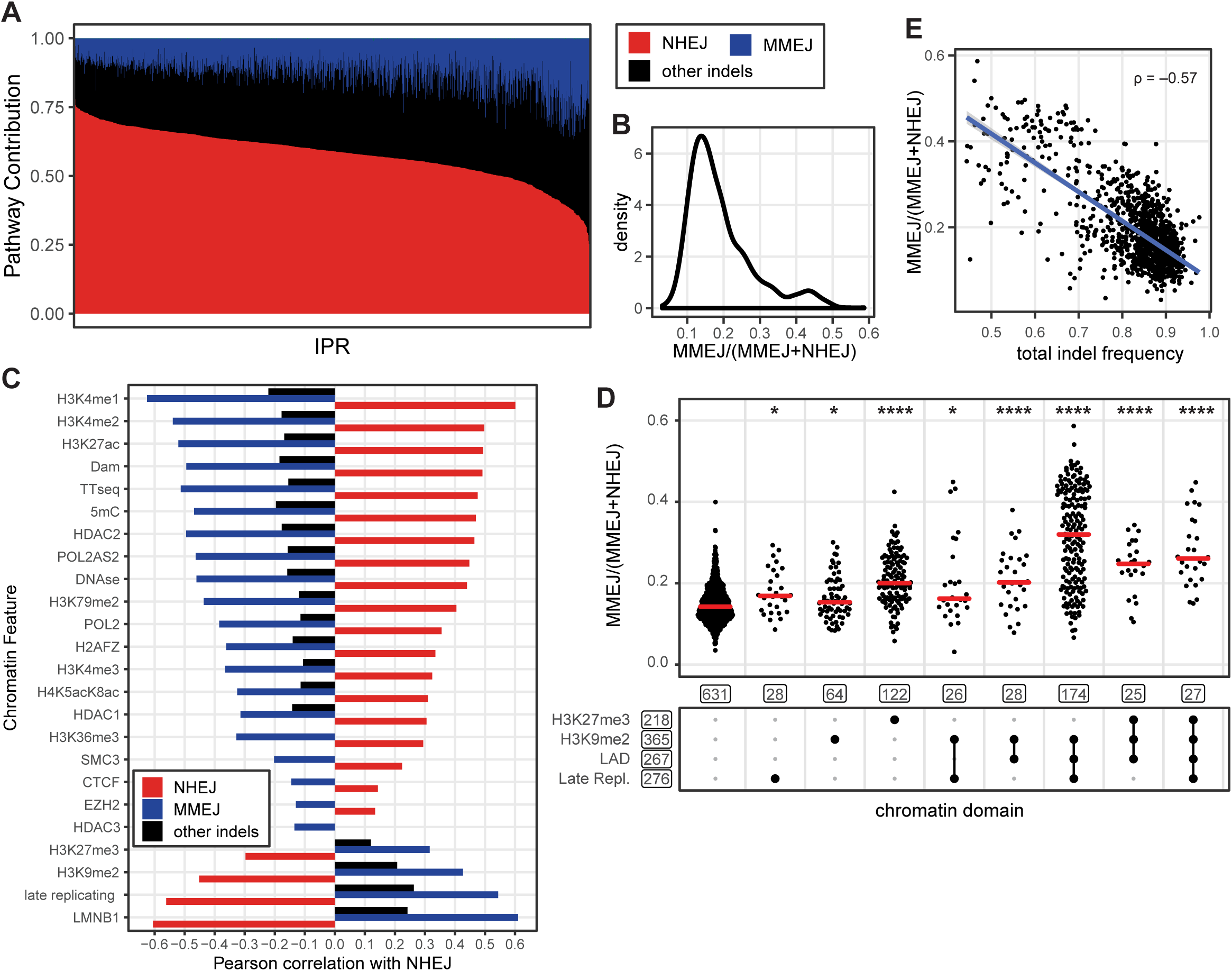
Different balance of NHEJ and MMEJ in LADs compared to inter-LADs. **(A)** Variation in indel composition across IPRs. Red: +1 (NHEJ); blue: −7 (MMEJ); black: other indels (unknown pathway). IPRs are ordered by +1 insertion frequency (1171 IPRs, 2-8 independent experiments). (**B**) MMEJ:NHEJ balance distribution across all IPRs. (**C)** Pearson’s correlation coefficients of the local intensities of 24 chromatin features versus the proportion of +1 (red), −7 (blue) and other indels (black). Correlations with p > 0.001 not shown. (**D**) MMEJ:NHEJ balance per IPR, split into different combinations of heterochromatin features as indicated by black dots in the scheme below the graph. Boxed numbers indicate the number of IPRs in each group; only groups with >20 IPRs are shown; see **Figure S4C** for all groups. Asterisks mark p-values according to the Wilcoxon test, compared to euchromatin IPRs (most left column): * p < 0.05, ** p < 0.01, *** p < 0.001, **** p < 0.0001. (**E**) Correlation between total indel frequency and MMEJ:NHEJ balance across all IPRs. Rho, Spearman’s rank correlation coefficient.

Overall, the patterns of MMEJ:NHEJ balance (**Figure 4D**) and indel frequencies (**Figure 3E**) appeared to mirror each other. Indeed, across all IPRs there is a substantial inverse correlation between both variables (**Figure 4E**). In other words, the shift towards MMEJ in heterochromatin is tightly linked to a reduction in total indel frequencies. It is likely that these reduced indel frequencies in heterochromatin are at least in part explained by lower cutting frequencies by Cas9, due to the compacted state of heterochromatin. However, it seems improbable that the cutting rate *itself* determines the pathway balance; possible models will be discussed below.

Other indels that could not be assigned to either MMEJ or NHEJ showed only weak correlations, with modest trends in the same direction as MMEJ (**Figure 4C**; **S4B**). Some of these larger deletions also tend to contain microhomologies but are more complex and too infrequent to be reliably categorized as MMEJ. Altogether, these data show that the balance between MMEJ and NHEJ is broadly linked to the global heterochromatin/euchromatin dichotomy, and within heterochromatin depends on the combination of heterochromatin features that are locally present.

### Gradual activation of MMEJ across chromatin types

To explore how the difference in pathway balance between heterochromatin and euchromatin develops over time after DSB induction, we conducted time series experiments. We used a robotics setup to collect DNA samples every three hours over a period of three days following Cas9 activation. For these experiments we focused on clone 5; because all 19 IPRs in this clone are in the same cell, their repair kinetics can be directly compared.

As expected, Cas9 activation resulted in a gradual accumulation of +1 and −7 indels in all IPRs, concomitant with a loss of wild-type sequence (**Figure 5A, S5**). These kinetics were generally slower in regions in LADs that are also marked by H3K9me2 and late replication (e.g. IPR 7, 14 and 16, **Figure S5**). Remarkably, the MMEJ:NHEJ balance was not constant over time, but was strongly skewed towards NHEJ at the early time points and gradually shifted towards MMEJ for all IPRs, culminating in a plateau approximately 50 hours after Cas9 activation (**Figure 5B**). This points to a slow activation of the MMEJ pathway, as we had observed previously for a single locus (Brinkman et al., 2018). This activation of MMEJ may occur eventually at most DSBs. However, over time the pathway balance diverged between IPRs in different chromatin environments, with several heterochromatic IPRs developing a higher MMEJ:NHEJ balance than most euchromatin regions (**Figure 5B**). This underscores that heterochromatic DSBs are intrinsically more prone to be repaired by MMEJ than euchromatic DSBs, but only once MMEJ is activated.

**Figure 5:**
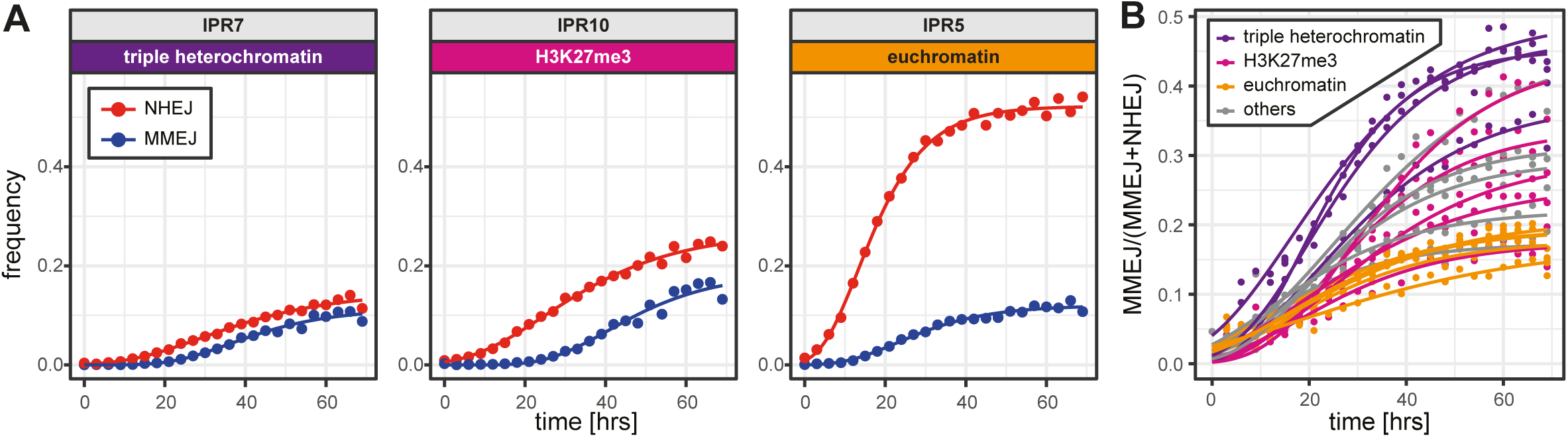
Accumulation of indels over time. (**A**) Time curves of the +1 insertion (red) and −7 deletion (blue) in single IPRs located in three different types of chromatin. See **Figure S5** for plots of all 19 IPRs in clone 5. Dots are measured values; lines are fitted sigmoid curves. Triple heterochromatin is the combination of H3K9me2, lamina association and late replication; euchromatin is here defined as the absence of any heterochromatin features. (**B**) Shifting MMEJ:NHEJ balance over time in 19 IPRs of clone 5. Triple heterochromatin IPRs in purple, H3K27me3 IPRs in magenta, euchromatin IPRs in orange and the other IPRs in gray. Data in A-B are averages of two independent experiments.

### Overall robustness of pathway balance in heterochromatin

We then investigated the role of several heterochromatin features in pathway balance by perturbation experiments. To distinguish direct from indirect effects, we compared the MMEJ:NHEJ balance of IPRs in regions with the targeted feature to that of IPRs in regions already lacking the feature prior to treatment. Direct effects should primarily alter the MMEJ:NHEJ balance in regions originally marked by the feature.

We first reduced the levels of H3K9me2 by treatment with the G9a-inhibitor BIX01294 (**Figure S6A-C**). This did not alter the MMEJ:NHEJ balance in H3K9me2 domains, except when in combination with LADs and late replication, where the balance increased slightly (p = 0.02; **Figure S6A**). We then tested the effect of GSK126, a compound that inhibits the H3K27me3 methyltransferase EZH2 and causes a global loss of H3K27me3 (**Figure S6D, E**). This inhibitor caused a significant reduction of the MMEJ:NHEJ balance in H3K27me3-only domains compared to euchromatin regions (p = 2e-11), as well as in virtually all domain combinations that were also covered by H3K27me3 (**Figure 6A**). Unexpectedly, GSK126 treatment also reduced levels of H3K9me2 (**Figure S6B, C**), and also slightly lowered the MMEJ:NHEJ balance in the triple H3K9me2, LAD and late replicating domains (p = 5.6e-05), but not in the single H3K9me2 domains. The most prominent shift in balance was, however, in H3K27me3 domains, pointing to a local effect of this histone modification on MMEJ:NHEJ balance.

**Figure 6:**
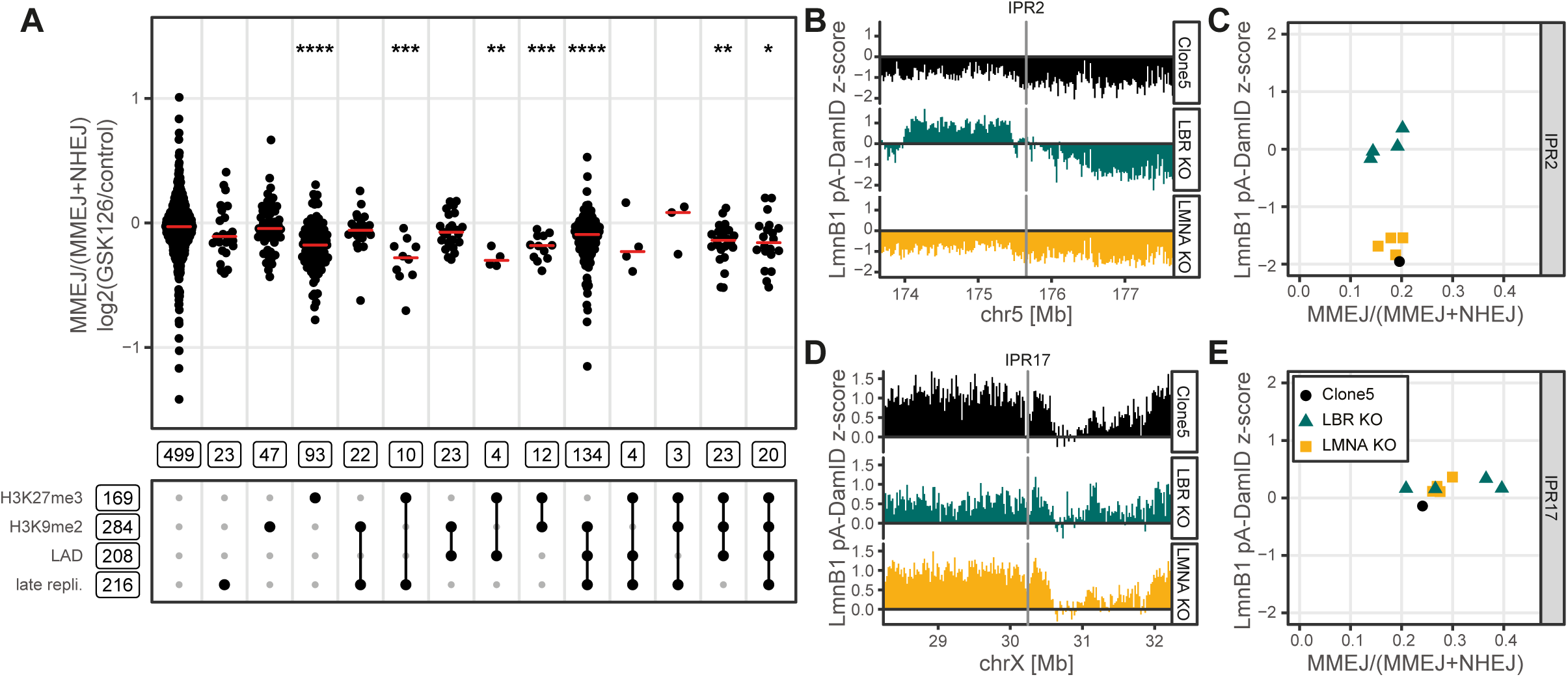
Effects of heterochromatin perturbations on pathway balance. (**A**) Log_2_ fold-change of MMEJ:NHEJ balance in GSK126 treated cells compared to control cells, for 917 IPRs divided by heterochromatin type. Data are average of two independent biological replicates. Wilcoxon test compared to euchromatin IPRs (most left column), * p < 0.05, ** p < 0.01, *** p < 0.001, **** p < 0.0001. (**B, D**) Nuclear lamina interaction tracks around IPR2 (**B**) and IPR17 (**D**). The tracks for the KO clones are average of 4 separate clones (individual tracks are shown in **Figure S7B**). All data are average of two independent biological replicates. (**C, E**) Comparison of MMEJ:NHEJ balance (n = 3) and average lamina interaction score in a 20 kb window centered on the IPR (n = 2), for IPR2 (**C**) and IPR17 (**E**).

Finally, because IPRs in LADs often show a high MMEJ:NHEJ balance, we used CRISPR/Cas9 editing to derive cell lines from clone 5 that lacked Lamin A/C (LMNA) or Lamin B Receptor (LBR) (**Figure S6F-I**). These two lamina proteins are important for the peripheral positioning of heterochromatin (Clowney et al., 2012; Solovei et al., 2013), and LMNA has been implicated in the control of NHEJ by sequestering 53BP1 (Redwood et al., 2011). Using the pA-DamID method (van Schaik et al., 2019) we mapped genome-wide changes in lamina interactions in four knock out (KO) clones each of LMNA and LBR. LMNA KO cells showed very few changes in lamina interactions, while the LBR KO clones showed a large number of regions with either gains or losses in lamina interactions. A detailed analysis of these changes will be reported elsewhere. Here, we investigated whether changes in lamina interactions of the IPRs coincided with changes MMEJ:NHEJ balance.

The majority of the IPRs did not undergo substantial changes in lamina interactions in either the LMNA or LBR KO clones compared to the parental clone 5, and they also did not show significant changes in MMEJ:NHEJ balance (**Figure S7A**). An exception was IPR2, in which the lamina interactions became stronger in all four LBR KO clones (**Figure 6B, C**). However, the MMEJ:NHEJ balance in IPR2 was not detectably altered in these clones (**Figure 6C, S7B**). This suggests that lamina contacts do not modulate this balance, but we cannot rule out that effects on this balance only emerge when lamina contacts are stronger than those of IPR2 in the LBR KO clones (note that the lamina interaction z-score in these clones is around 0, which corresponds to the genome-wide average level of lamina interactions). Interestingly, IPR17 showed a marked increase in the MMEJ:NHEJ balance in two of the four LBR KO clones (**Figure 6D-E, S7A**). However, for this IPR the lamina interactions did not change (**Figure S7C**), and we do not understand why only two of the four clones show this behavior. Nevertheless, this result underscores that it is possible to shift the MMEJ:NHEJ balance in an IPR markedly without any change in its sequence. Presumably an unknown change in the local chromatin state in the two clones is responsible for this.

Together, these data indicate that the MMEJ:NHEJ balance in specific heterochromatin types is not easily shifted by targeting individual key markers of the respective heterochromatin types. Pathway balance may be redundantly controlled by multiple factors in each heterochromatin type. Nevertheless, depletion of H3K27me3 did cause a detectable reduction in MMEJ:NHEJ balance in heterochromatin domains that normally carry this mark.

### Impact of chromatin context on SSTR

Finally, we investigated a third repair pathway: SSTR, which is commonly used to create specific mutations by CRISPR/Cas9 editing. We hypothesized that this pathway competes with NHEJ and MMEJ, and that its relative activity may also be modulated by the local chromatin environment. To test this, we triggered DSB formation in our reporter sequence in the presence of a template ssODN containing a specific +2 insertion (ssODN insertion) (**Figure S8A**). We designed this insertion within the PAM site, so that a successful editing event destroyed the PAM site and prevented further cutting by Cas9. We then transfected the IPR cell pools with this ssODN (together with the sgRNA) to probe the impact of chromatin context on the relative activity of SSTR, NHEJ and MMEJ in parallel. In these experiments, we found that a median 6% of the indels consisted of the SSTR insertion (**Figure S8B**). Accumulation of this insertion was mostly at the expense of the −7 deletion but not of the +1 insertion (**Figure 7A, S8C, D**), suggesting a competition between SSTR and MMEJ. Indeed, depletion of POLQ (a key protein of MMEJ) caused an increase in ssODN insertions (**Figure S2D, E**). Consistent with an earlier study (Richardson et al., 2018), we find that knockdown of CtIP, a factor that promotes end-resection (Sartori et al., 2007) strongly reduces SSTR and MMEJ activity (**Figure S2D, E**). This suggests that DSB end-resection is an early step in SSTR. Furthermore, in the presence of DNA-PKcs inhibitor NU7441 the ssODN insertion was increased while the +1 insertion was reduced (**Figure 7A, Figure S2E**), indicating that the ssODN insertion does not result from NHEJ. In agreement with previous work (Richardson et al., 2018) we conclude that SSTR is distinct from NHEJ and MMEJ.

**Figure 7:**
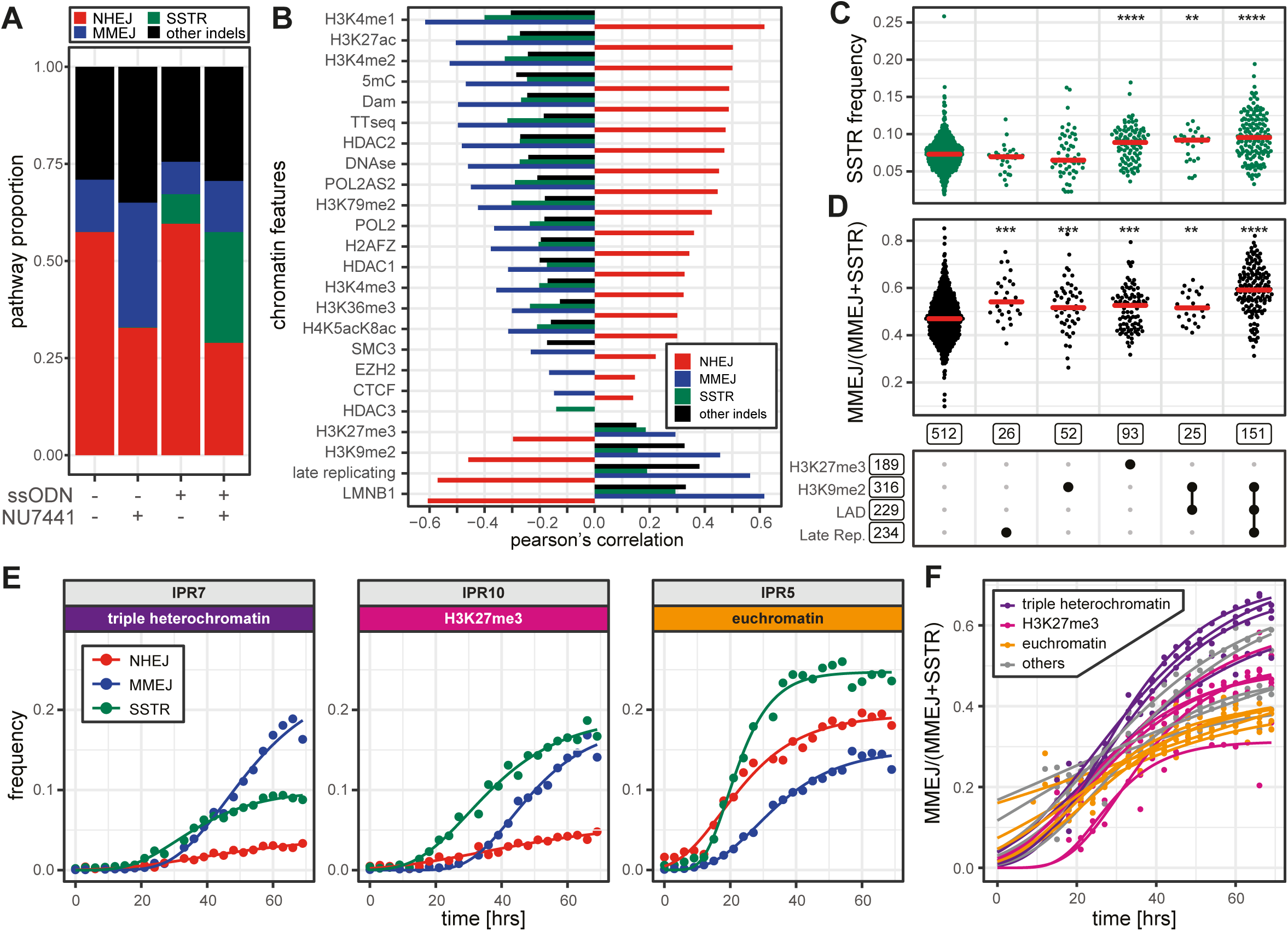
Balance between NHEJ, MMEJ and SSTR in different chromatin contexts. (**A**) Average pathway contribution across all IPRs in the cell pools, in the absence or presence of a ssODN donor, and with or without NU7441 treatment (n = 2-8). Red: +1 insertion (NHEJ); blue: −7 deletion (MMEJ); green: +2 insertion due to SSTR; black: other indels. (**B**) Pearson’s correlations of the local intensities of 24 chromatin features versus the relative activity of each pathway (n = 2-3). Correlations with p > 0.001 not shown. Colors as in **A**. (**C**) Proportion of DSBs repaired by SSTR in the presence of the ssODN donor, for each IPR, split according to the heterochromatin features present as indicated by black dots in the scheme below the graph. Boxed numbers indicate the number of IPRs in each group; only groups with >20 IPRs are shown; see **Figure S8I** for all groups. Asterisks mark p-values according to the Wilcoxon test, compared to euchromatin IPRs (most left column): * p < 0.05, ** p < 0.01, *** p < 0.001, **** p < 0.0001. (**D**) Same as in C, but showing the MMEJ:SSTR balance. See also **Figure S8J**. (**E**) Time curves as in Figure 5A, but now in the presence of the ssODN donor and NU7441 (n = 2). Colors as in **A**. Dots are measured values; lines are fitted sigmoid curves. See **Figure S9** for plots of all 19 IPRs in clone 5. (**F**) Gradual increase of the MMEJ:SSTR balance over time in 19 IPRs of clone 5. Triple heterochromatin IPRs in purple, H3K27me3 IPRs in magenta, euchromatin IPRs in orange and the other IPRs in gray.

MMEJ and SSTR correlate rather poorly with each other across the IPRs (**Figure S8E**). In the presence of NU7441 this correlation was even weaker (**Figure S8F**). This lack of correlation implies that the two pathways are largely subject to different local control mechanisms. We therefore searched for chromatin features that may explain this. Like MMEJ, SSTR generally correlates positively with heterochromatin features and negatively with euchromatin features, but the correlations are overall weaker than observed for MMEJ and NHEJ (**Figure 7B; S8G, H**). The proportion of DSBs repaired by SSTR is highest in heterochromatic regions marked by H3K9me2 and LADs, either with or without late replication; and in regions marked by H3K27me3 (**Figures 7C, S8I**). Searching for chromatin features that might explain the MMEJ:SSTR balance differences, we found only minor differences in this balance between chromatin types, with regions triple-marked by H3K9me2, LADs and late replication showing the most pronounced skew towards MMEJ. Other regions marked by one or two heterochromatin features showed only marginal effects compared to euchromatic regions (**Figure 7D, S8J**).

Finally, we compared the timing of SSTR and MMEJ. For this, we conducted time series experiments in the presence of NU7441 to reduce NHEJ activity, in order to maximize barcode reads with SSTR and MMEJ indels (**Figure 7E, S9**). Individual time curves suggest that SSTR tends to become active at the DSBs earlier than MMEJ. Indeed, plotting the MMEJ:SSTR balance as a function of time shows that this balance is initially strongly skewed towards SSTR, and only gradually shifts towards MMEJ (**Figure 7F**). This shift is particularly pronounced in heterochromatin triple-marked by H3K9me2, LADs and late replication, and slightly less in H3K27me3 regions (**Figure 7F**). Altogether these results demonstrate that MMEJ and SSTR compete, that their balance is at least in part controlled by the local chromatin context, and that SSTR is activated earlier than MMEJ.

## DISCUSSION

Here, we present a powerful reporter system to query effects of chromatin on DSB pathway usage. It consists of (1) a simple short DNA sequence that, when cut with Cas9, produces a signature indel for each repair pathway; and (2) an adaptation of the TRIP multiplexed reporter assay (Akhtar et al., 2013). In combination, these tools offer precise measurements of the relative activity of NHEJ, MMEJ and SSTR, combined with the throughput that is needed to query the impact of a wide diversity of chromatin contexts. The sequencing-based “scar-counting” readout renders the assay highly quantitative and can provide detailed insights into the repair kinetics. Moreover, the large number of randomly integrated IPRs provides a broad sampling of all common chromatin contexts (Akhtar et al., 2013).

Using this approach in human cells, we found that the balance between MMEJ and NHEJ varies >5-fold across chromatin contexts. Generally, heterochromatin is more prone to repair by MMEJ than euchromatin, but this shift depends on the precise heterochromatin features that are present. It is possible that euchromatin carries one or more features that activate or recruit the NHEJ machinery, or conversely that certain heterochromatin features promote MMEJ. If the latter is true, then it should be considered that multiple heterochromatin features can play such a role, since H3K27me3 modulates the MMEJ:NHEJ balance in H3K27me3-marked heterochromatin, but this does not explain the high MMEJ:NHEJ balance in heterochromatin marked by late replication, H3K9me2 and lamina interactions.

An alternative model builds on our observation that MMEJ is only slowly activated. A DSB in “open” euchromatin may be rapidly accessed and repaired by NHEJ, often before MMEJ has been activated. In contrast, in heterochromatin a DSB may be inaccessible to either pathway until the heterochromatin is de-compacted. This remodeling of heterochromatin may be a slow process, which would allow time for MMEJ to be activated and giving both MMEJ and NHEJ a more similar chance to repair the DSB. Indeed, DSB-induced unfolding of heterochromatin has been reported (Goodarzi et al., 2008; Chiolo et al., 2011; Jakob et al., 2011; Ryu et al., 2015; Janssen et al., 2016; Tsouroula et al., 2016). However, as seen by microscopy, this remodeling process occurs within ∼20 minutes, while we found that MMEJ activation takes several hours. It is possible that additional biochemical or structural changes in heterochromatin are involved that take place over a time scale of hours. We also considered that one early cutting event in euchromatin may trigger slow upregulation of MMEJ activity globally throughout the nucleus, which would then increase the probability of MMEJ repairing a DSB that is formed later in heterochromatin (which may be cut more slowly). However, this explanation seems unlikely, because early breaks caused by ionizing radiation do not boost MMEJ repair at a Cas9 cut ∼16 hours later (Brinkman et al., 2018). This result suggests that the slow MMEJ activation does not occur globally throughout the nucleus, but rather locally at the DSB.

Interestingly, SSTR does not show the same delay as MMEJ, and has a clear advantage over MMEJ in heterochromatin early after DSB induction. Because both pathways require end-resection, this result implies that end-resection is not the rate limiting step responsible for the delayed MMEJ. Indeed, rapid localization of end-resection proteins at DSBs inside heterochromatin has been observed (Chiolo et al., 2011).

As summarized in the Introduction, previous studies have addressed the impact of specific chromatin contexts or proteins on NHEJ and HR, but generally did not monitor MMEJ and SSTR. While the latter two pathways share components with HR, they are mechanistically distinct, and thus the effects of chromatin context may differ. For example, several previous studies have indicated that H3K36me3, which is generally present along active transcription units, promotes HR (Aymard et al., 2014; Carvalho et al., 2014; Pfister et al., 2014; Clouaire and Legube, 2015). We find H3K36me3 to correlate negatively with MMEJ. Possibly, HR and MMEJ respond differently to H3K36me3, or the repair of Cas9-induced breaks differs from breaks induced by other means.

The activities of MMEJ and SSTR that we detect indicate that end-resection is generally not impeded by various types of heterochromatin. Previous work has pointed to a role of HP1α and HP1β in recruiting proteins involved in end-resection (Soria and Almouzni, 2013), but not all types of heterochromatin are marked by these proteins. Our data indicate that multiple heterochromatin components contribute locally to promoting MMEJ and SSTR. This includes LADs, which is in agreement with a previous study that implicated various MMEJ-specific proteins in the repair of DSBs near the nuclear lamina (Lemaitre et al., 2014).

The results obtained in this study have practical implications for genome editing by means of Cas9. First, the efficiency of Cas9 editing is generally lower in most types of heterochromatin compared to euchromatin. This has been noted before, but based on data that covered only a small number of loci that did not compare all heterochromatin types (Chen et al., 2016; Daer et al., 2017; Jensen et al., 2017; Kallimasioti-Pazi et al., 2018). Our data indicate that Cas9 editing is primarily suppressed in regions that carry a combination of lamina interactions, late replication and H3K9me2. Most likely the relatively low accessibility of the DNA in these loci is preventing efficient cutting by Cas9. Regions that carry only one of these marks, or H3K27me3, show only a modestly reduced editing efficiency.

From a genome editing perspective, the skew towards the MMEJ and SSTR pathways in heterochromatin is a convenient compensation for the lower overall editing efficiency, because MMEJ and SSTR are generally more useful than NHEJ to generate specific types of mutations. MMEJ is better suited to generate frameshifts and deletions that can result in functional knockout of genes, while SSTR is particularly useful to generate specifically designed mutations. Maps of heterochromatin features are thus useful resources to choose the optimal target loci for CRISPR/Cas9 editing, particularly when combined with algorithms that predict editing outcomes based on sequence (van Overbeek et al., 2016; Allen et al., 2018; Shen et al., 2018; Chakrabarti et al., 2019; Chen et al., 2019).

This work complements a recent study that employed a multiplexed reporter for DNA mismatch repair, which did not find significant effects of chromatin context on the repair outcome (Pokusaeva et al., 2019). Another multiplexed integrated reporter study also found evidence that genomic location can affect Cas9 editing efficiency (Gisler et al., 2019), but these results were more difficult to interpret because the reporter sequence itself was not transcriptionally inert. Importantly, neither of these studies addressed the impact of chromatin context on the balance between specific DSB repair pathways. Our multiplexed reporter assay provides new opportunities to systematically investigate the role of local chromatin context in DSB repair by multiple pathways. Moreover, our time-series experiments demonstrate that the assay can be performed in 96-well format, making it scalable for applications such as drug screens and CRISPR screens. In the future, the assay may also be modified to include the detection of DSB resection or other intermediates, and perhaps of HR activity.

## Acknowledgments

We thank the NKI Genomics, Flow Cytometry and Research High Performance Computing core facilities for excellent support, and members from our laboratories for inspiring and helpful discussions. Supported by ZonMW-TOP grant 91215067 (to R.H.M. and B.v.S.) and NIH Common Fund “4D Nucleome” Program grant U54DK107965 (B.v.S.). SGM is funded by Marie-Curie/AIRC iCARE2.0 fellowship 800924. The Oncode Institute is partly supported by KWF Dutch Cancer Society.

## Author contributions

Conceived and designed study: E.K.B., R.S., R.H.M., B.v.S.

Experiments: R.S., E.K.B., X.V., T.v.S., S.G.M., D.P.H., B.M., J.v.d.B.

Data processing, bioinformatics, figure preparation: C.L., R.S., E.K.B., T.v.S., X.V., S.G.M.

Supervision, project management: B.v.S., R.H.M., R.L.B.

Manuscript writing: R.S., E.K.B., B.v.S., with input from all authors.

## METHODS

### Constructs

The IPR-PB-BC (PiggyBac) construct was derived from the pPTK-Gal4-tet-Off-Puro-IRES-eGFP-sNRP- pA-trim1 plasmid (GenBank accession KC710229). The enhanced green fluorescent protein (eGFP) expression transcription unit including promoter, the puromycin resistance cassette (PuroR) and the internal ribosome entry site (IRES) were replaced with the sgRNA-LBR2 target sequence and its flanking region. This plasmid did contain a point mutation which was removed by restriction cloning using a derivative of plasmid with a shortened 3’ITR of 67 bp (pPTK-P.CMV.584-eGFP-trim1-PI04 – kindly provided by Alexey Pindyurin and Waseem Akhtar). The target sequence was obtained by annealing ODS001 and ODS002 (400 pM each) (for primer sequences see **Table S4**) in 50 µl MyTaq Red mix followed by 5 cycles of PCR. This PCR product was then further amplified with TAC0001 and TAC0002 (50 pM each). This sequence was then inserted in the PB backbone by restriction cloning with NheI and KpnI. This construct (IPR-PB) was then used to make the barcoded plasmid libraries, the 3′-ITR of PiggyBac was amplified with primers TAC0003 (containing a 16 nucleotide random barcode) and TAC0004. The PCR product was digested with KpnI and BssHII, ligated into the KpnI and MluI sites of the IPR-PB plasmid and transformed into CloneCatcher DH5α electrocompetent *E. coli* (Genlantis - C810111). A pool of ∼500,000 transformed bacterial cells were grown and plasmids were purified, resulting in the IPR-PB plasmid library. The PB transposase expression vector (mPB-L3-ERT2-mCherry) is described in (Akhtar et al., 2014). The sgRNAs were designed using CHOPCHOP (Montague et al., 2014) and cloned into expression vector pBlue-sgRNA (Brinkman et al., 2018) (see **Table S4**). The sgRNA sequences are listed in **Table S3**.

### Cell culture

We used clonal cell line K562#17, which is a human K562 cell line stably expressing DD-Cas9 (Brinkman et al., 2018). K562#17 cells were cultured in RPMI 1640 (Life Technologies) supplemented with 10% fetal bovine serum (FBS, HyClone®), 1% penicillin/streptomycin. Mycoplasma tests, performed every 1-2 months, were negative.

### Generation of IPR cell pools

Cell pools carrying IPRs were produced as described (Akhtar et al., 2014). Briefly, K562#17 cells were transfected with 32 μg of barcoded IPR-PB-BC plasmid library and 6 μg of PB transposase plasmid using Lipofectamine 2000 (ThermoFisher #11668019). Mock-transfected (without PB transposase) and GFP plasmid controls were included. After 24 h, the cells were sorted by fluorescence-activated cell sorting (FACS) based on mCherry signals. We discarded cells without any detectable mCherry signal, because they most likely failed to take up any plasmid. 0.5 μM of 4-hydroxytamoxifen (4-OHT) was added to the samples to activate the transposase. Sixteen hours later the cells were washed to remove 4-OHT. After sorting, the population was grown for 8 days to clear the cells from free plasmid. Then, the mCherry negative cells were FACS sorted in aliquots of ∼2000 cells, which were expanded to establish two cell pools, each with a different collection of IPRs. We also isolated single cells to make clonal TRIP lines, including clone 5.

### LMNA & LBR knock out generation

One million clone 5 cells were transfected with 3 µg of plasmid expressing the following sgRNAs per clone: LMNA_KO1 & LMNA_KO2 (each 1.5 µg) for LMNA KO 1 and 2; LMNA_KO4 for LMNA KO 3 and 4; LBR_KO1 for LBR KO 1-4 (**Table S3**), and cultured in complete RPMI medium with 500 nM Shield-1 (to activate Cas9) for 3 days. To obtain individual clones, cells were plated in two 96-well plates by limiting dilution (2 cells per mL; 100 µl per well). Each clone was then tested by TIDE (Brinkman et al., 2014) for frameshifts in all alleles (primers in **Table S3**). For each sgRNA we selected two clones with complete frameshifts for further experiments.

### siRNA lipofection

All siRNAs were obtained from Dharmacon as ON-TARGETplus Smartpool and transfected with the RNAiMAX Transfection kit (Thermo) at a final concentration of 25 nM, 24h prior to sgRNA electroporation. Samples were collected 24h after electroporation for subsequent western blotting analysis.

### Western blots

Whole-cell extracts of ∼0.5×10_6_ cells were prepared by washing cultures in PBS and lysing with 50 μL lysis buffer (Tris pH 7.6, 10% SDS, Roche cOmplete™ Protease Inhibitor Cocktail). Western blotting was performed according to standard procedures using the following antibodies and dilutions: H3K27me3 (1:1000 Cell Signaling C36B11, rabbit), H3K9me2 (1:1000 Upstate 07-441, rabbit).

### Transfection of sgRNA plasmids and ssODN

For transient transfection of the sgRNAs, 1 to 6 ×10_6_ cells (lower limit for clonal experiments, higher limit for pooled experiments) were resuspended in transfection buffer (100 mM KH_2_PO_4_, 15 mM NaHCO_3_, 12 mM MgCl_2_, 8 mM ATP, 2 mM glucose (pH 7.4)) (Hendel et al., 2014). After addition of 3.0-9.0 µg plasmid, the cells were electroporated in an Amaxa 2D Nucleofector using program T-016. DD-Cas9 was induced directly or ∼16 hours after transfection with a final concentration of 500 nM Shield-1 (Aobious). To probe SSTR, 3-9 µg sgRNA was co-transfected with 1.5-4.5 µg ssODN (5’ TAGAATGCTAGCGTGACTGGAGTTCAGACGTGTGCTCTTCCGATCTAATTTCTACTTCATAAT AAAGTGAACTCCCAGGCCATCGAC<u>AT</u>CTCTTACCACTTCACCATCGGCAAATTTCCTACTTG GCATT 3’, Ultramer grade, IDT). The specific mutation that disrupts the PAM is underlined.

### Inhibitor treatments

DNA-PKcs inhibitor NU7441 (Cayman; diluted 1:1000 from 1 mM stock in dimethylsulfoxide [DMSO]), M3814 (MedChemExpress; diluted 1:1000 from 1 mM stock in DMSO), GSK126 (Selleckchem; diluted 1:2000 from 1 mM stock in DMSO), BIX01294 (Sigma; diluted 1:1000 from 1 mM stock in H_2_O), or respective solvent-only controls at equal volumes, was added to the cells at the same time when the cells were supplemented with Shield-1 to induce DD-Cas9 or 24 hours prior to nucleofection for GSK126 and BIX01294. DMSO was also present in the experiments in figures 3, 4, 5, 6b, 7, S2, S4, S5, S6a-d, S7.

### TIDE method

The TIDE method was performed as described in (Brinkman et al., 2014). Briefly, PCR reactions were carried out with ∼100 ng genomic DNA in MyTaq Red mix (Bioline) and purified using the PCR Isolate II PCR and Gel Kit (Bioline) or by ExoSAP (for primers see **Table S4**). ExoSAP was done by adding 0.125 µl Shrimp Alkaline Phosphatase (1U/µl; New England Biolabs, M0371S), 0.0125 µl Exonuclease I (20 U/µl; New England Biolabs, M0293S) and 2.3625 µl H_2_O per 10 µl PCR reaction. Samples were incubated 30 minutes at 37 °C and inactivated for 10 minutes at 95 °C. About 2 µl (50-100 ng) of purified PCR product was then subjected to Sanger Sequencing by Eurofins Genomics. The sequence traces were analyzed using the TIDE analysis tool (https://tide.nki.nl).

### Immunostaining of the KO clones

Coverslips were first coated with 0.1% (w/v) poly-L-lysine (Sigma-Aldrich, #P8920) for 15 minutes, washed with H_2_O (1x) and PBS (3x) and stored in 70% ethanol for later use. 1 × 10_6_ cells for each KO clone of LMNA and LBR were collected, centrifuged (3 minutes, 500 g) and washed once in PBS. Cells were resuspended in 90 μl PBS and added dropwise to a dry, poly-lysine coated coverslip (for WT 300 μl PBS over 4 coverslips). After 10 minutes, cells were fixed by adding dropwise 1 ml of 2.5% formaldehyde in PBS and fixed for 10 min at room temperature (RT). Coverslips were washed once with PBS and permeabilized cells with 0.5% NP40/PBS for 10 min at RT and washed again with PBS. Coverslips were transferred cell-side up on parafilm in a hybridization container and blocked for 15 minutes at RT in 90 μl 1% BSA/PBS, before overnight incubation at 4°C with primary antibody mixes (1:500 Lamin B1 antibody (Abcam ab16048, rabbit), 1:200 Lamin B2 antibody (Abcam ab8983, mouse), 1:100 LBR antibody (Abcam ab122919, rabbit) or 1:100 Lamin A/C antibody (Cell Signaling sc-6215, goat)). After three washes with 1% BSA/PBS, coverslips were incubated with a secondary antibody. For Lamin B1 (rabbit) we used 1:100 B26 Jackson rabbit 594 (Jackson 711-585-152). For LaminB2 (mouse) we used 1:100 B25 Jackson mouse 594 (Jackson 715-585-150). For LBR (rabbit) we used 1:100 B12 rabbit FITC (Jackson 711-095-152). For Lamin A/C (goat) we used 1:100 B13 goat FITC (Jackson 705-095-147). This was followed by washes with PBS (3x) and H_2_O (1x), and mounted with Vectashield + DAPI (Vector Laboratories, #H-1200).

### pA-DamID

pA-DamID maps were generated and processed as described (van Schaik et al., 2019). Briefly, 1 million cells were collected by centrifugation (3 minutes, 500g) and washed in ice-cold PBS and subsequently in ice-cold digitonin wash buffer (DigWash) (20 mM HEPES-KOH pH 7.5, 150 mM NaCl, 0.5 mM spermidine, 0.02% digitonin, protease inhibitor cocktail). Cells were resuspended in 200 µL DigWash with 1:100 mouse Lamin B2 antibody (Abcam, ab8983) and rotated for 2 hours at 4°C, followed by a wash step with 0.5 mL DigWash buffer. This was repeated with a 1:100 mouse anti-rabbit antibody (Abcam, ab6709) and 1 hour of rotation, and afterwards with 1:100 pA-Dam (∼60 NEB units). After two washes with DigWash, cells were resuspended in 100 µL DigWash supplemented with 80 µM SAM to activate Dam and incubated for 30 minutes at 37°C. Genomic DNA was extracted using the ISOLATE II Genomic DNA kit (Bioline cat. no. BIO-52067) and DNA was processed for high-throughput sequencing similar to conventual DamID (Vogel et al., 2007; Leemans et al., 2019), except that the DpnII digestion was omitted. To control for DNA accessibility and amplification bias, 1 million permeabilized cells (without any antibodies bound) were incubated with 4 units of Dam enzyme (NEB, M0222L) during the activation step. This sample functions as “dam-control” over which a log2-ratio is determined. Log2-ratios were converted to z-scores to account for small differences in dynamic range between experiments.

### Generation of indel sequencing libraries

After 64 hour incubation, the cells were collected and genomic DNA was extracted using the ISOLATE II Genomic DNA kit (Bioline cat. no. BIO-52067). PCR was performed in two steps and pooled experiments were performed in triplicates for a higher coverage. IndelPCR1 was performed with 200 ng genomic DNA each using primers TAC0007 (indexed) and TAC0012 that amplify 1 bp upstream of the barcode 46 bp downstream of the cut-site (see **Figure 1A, Table S4**). indelPCR2 used 2 µl of each indelPCR1 product with TAC0009 and either TAC0011 (non-indexed) or TAC0159 (indexed). Each sample was generated with a unique combination of one or two indexes. Both PCR reactions were carried out with 25 µl MyTaq Red mix (Bioline cat. no. BIO-25044), 0.5 µM of each primer and 50 µl final volume. PCR conditions for both steps were 1 min at 95 °C, followed by 15 sec at 95 °C, 15 sec at 58 °C and 1 min at 72 °C (5x), followed by 15 sec at 95 °C, 15 sec at 65 °C and 1 min at 72 °C (10x). The indelPCR2 was pooled per experiment after quantification on a 1% agarose gel and cleaned up using CleanPCR (CleanNA) beads at 0.8:1 beads : sample ratio. The purified PCR product was run on a 1% agarose gel cut out to remove remaining primer dimers and cleaned with PCR Isolate II PCR and Gel Kit (Bioline). The purified libraries were sequenced on an Illumina HiSeq2500 or MiSeq depending on the expected complexity of the library.

### Time Series

Sample collection in the time series experiments was done automatically using a Hamilton Microlab® STAR equipped with a Cytomat 2 C450 incubator. One million cells were transfected as described above with either the sgLBR2 plasmid alone or with the ssODN. After 16 hours, 40,000 cells in 100 µl medium were seeded per well in 96-well plates. The automated system then added 100 µl RPMI medium (with 1 µM Shield-1) to each well, for a final 500 nM Shield-1 concentration. The first time point was directly collected and required a brief centrifugation step (10 seconds at 300g) to precipitate the cells, before returning the cell culture plate to the robot. Then for each timepoint 170 µl medium was removed from the well and discarded, the left-over was mixed and transferred to a new 96-well PCR plate at 8 °C. Each newly collected well was then filled with 50 µl of DirectPCR® Lysis (Viagen Cat. No. 302-C) buffer with 1 mg/ml Proteinase K (Bioline, Cat. No. BIO-37084) to pre-lyse the cells. The cell culture plate was returned to the incubator and every 3 hours a new timepoint was collected as described above. One 96-well plate included 4 timeseries of each 24 timepoints. After 69 hours the collection of samples was finished and the cell lysates were sealed and incubated for 3 hours at 55 °C and heat-inactivated for 10 minutes at 95 °C.

### Timeseries sequencing library preparation

Library preparation for the timeseries was very similar to the pool experiments except for indelPCR1. 20 µl of crude lysate was used in a total PCR volume of 80 µl, with 40 µl MyTaq HS Red mix (Bioline, BIO- 25048) and 0.5 µM of each primer. PCR cycles were as described above.

### Mapping of IPR integration sites: experimental methods

Mapping of IPR integration sites was performed in two replicates by inverse PCR (iPCR) followed by 2 x 75 bp paired end sequencing on an Illumina HiSeq2500 as previously described (Akhtar et al., 2014).

### Mapping of IPR integration sites: computational methods

Linking of IPR barcodes to the integration sites was adapted from (Akhtar et al., 2013). Reads of both replicates were pooled. The first read in each read pair was used to extract the barcode. This was done using the ‘GTCACAAGGGCCGGCCACAAC’ constant sequence followed by a regular expression ‘TCGAG[ACGT]{16}TGATC’. From the sequence matching this regular expression, the 16 bp barcode was extracted. To identify barcodes arising from mutations during PCR and sequencing, *starcode v1*.1 (Zorita et al., 2015) was used with the sphere clustering setting and a maximum Levenshtein distance of 2. The second read of each pair was used to locate the site of integration after removing the ‘GTACGTCACAATATGATTATCTTTCTAGGGTTAA’ sequence matching the transposon arm. The flanking sequence was aligned to GRCh38 using *bowtie2* using the *very-sensitive-local* option (20 seed extension attempts, up to 3 re-seed attempts for repetitive seeds, 0 mismatches per seed, with a seed-length of 20 and using a multi-seed function: 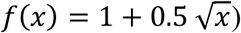. Locations of integration sites were required to be supported by at least 5 reads with an average mapping quality larger than 10 at the primary location, having at least 95% of the reads located at this locus, with not more than 2.5% of the reads at a secondary location.

### Indel scoring

Indel reads after induction and repair of DSBs consist of single-end reads of 150 bp that span both the DSB site and the barcode. Indel scoring was adapted from (Brinkman et al., 2018). Barcodes were extracted from the reads with an in-house script using functions of *cutadapt 1*.*11*. The 16 bp barcode was located using the 20 bp constant ‘GTCACAAGGGCCGGCCACAA’ sequence preceding the barcode and ‘TGATCGGT’ expected immediately after the barcode. For the 20 bp constant sequence, 2 mismatches were allowed.

To determine the indel size in each read, we used the distance (number of nucleotides) between two fixed sequences at the start and at the end of the read. The indel size was calculated as the difference between the measured distance and the expected distance based on the wild-type sequence. We used the following anchor sequences: before the break site, ‘TGATCGGT’ and after the break site, ‘GGAGTT’, ‘CACTTT’, ‘ATTATG’, ‘GAAGTA’ and ‘ATTAGA’. The most proximal match found with these sequences was used to calculate the indel size by subtracting the expected location from the observed location. Insertions and deletions have indel sizes >0 and <0, respectively. Wild-type sequence is defined as indel size 0. Point mutations were not analyzed. Per replicate experiment we observed a median 16.4% sequence reads in which we could not find a match with the constant parts; we discarded these reads in subsequent analyses. Potentially these represent large deletions, complex mutations, sequencing errors or a combination thereof.

Per barcode, the reads of all technical replicates were pooled if applicable. Mutated barcodes were included or discarded as described above for the mapping of IPR integrations. Because in the cell pools not all IPRs are equally represented (the cell pools consist of a mix of clones that each carry different IPRs, and some cell clones grow faster than others), we then discarded IPRs that were too underrepresented to provide reliable data. Specifically, we required that each IPR is represented by at least 50 cells among the ∼100,000 cells that were used in each experiment. We assumed an average of 6 IPRs per cell. Accordingly, the number of total reads per IPR was divided by the library size and multiplied by 6 * 100000 to obtain the estimated number of cells for each IPR. IPRs for which this score was >50 were used for subsequent analyses. Then each replicate was normalized over library size and biological replicates were averaged. The frequency of each indel type as proportion of total reads was calculated on that average. Pathway frequency per IPR was calculated as a proportion of the specific mutation over all indels (excluding wild-type sequences).

### Preprocessing of previously published epigenome data

Published ChIP-seq data from various sources (**Table S1**) were re-processed for consistency. Raw sequencing data were obtained from the Sequence Read Archive (https://www.ncbi.nlm.nih.gov/sra/). Reads were aligned to the human genome GRCh38 using *bowtie2* with default options. Replicate datasets were processed separately, while the sequences from the input were combined. After alignment, reads were filtered on a minimum mapping quality of 30. Duplicate reads were removed except for the reads coming from experiments using tagmentation, in which duplicates were kept. After this, genomic regions were masked based on blacklist regions identified by the ENCODE project (ENCFF419RSJ) (Encode Project Consortium, 2012) and putative artifact regions were identified based on the input reads using *chipseq-greylist*, a python implementation of GreyListChIPs (https://doi.org/doi:10.18129/B9.bioc.GreyListChIP). We considered ChIP-seq datasets to be of sufficient quality for our analyses if there was well annotated input and sample data available and consistent read lengths were used. Mean ChIP-seq signals for IPR integration sites were calculated by taking the sum of the reads in a region of 2 kb around the IPR, scaling input and sample counts by the smallest library size, adding a pseudo count of 1 and subsequently dividing sample over input normalized counts. After this, replicate experiments were averaged. For domain calling, ChIP-seq signals were calculated in similar fashion for bins of 5kb. *HMMt*, an R package implementing a Hidden Markov model with t emission, was used to subsequently call domains (https://github.com/gui11aume/HMMt).

DamID data of Lamin B1 are from (Leemans et al., 2019). The DamID score was calculated by scaling counts to the smallest library size, adding a psuedocount of 1 and dividing over Dam-only.. The normalized dam-only score was log_2_-transformed before averaging between replicates to calculate the dam accessibility score. Replication timing data was obtained from the 4DN data portal in the form of read coverage for late and early fraction separately. Counts were processed in the same way as for the ChIP data. For TTseq coverage from forward and reverse tracks were summed and the lowest coverage score above zero was used as pseudo count before log_2_-transforming and averaging between replicates. For DNAse hypersensitivity data of both paired-end and single-end sequencing reactions were used from encode. Coverage tracks were used and for the single ended reaction a small pseudo count of half the minimum value above 0 was used before log_2_ transforming. Paired-end and single-end coverage was log_2_ transformed before averaging. Whole genome bisulfite sequencing tracks from encode were used and coverage was calculated and log_2_ transformed without the need for a pseudo count. Replicates were subsequently averaged. Data sources are available in **Table S2**.

Z-scores of above chromatin information for the clonal line was calculated by using the mean and standard deviation of the signals in the TRIP pool.

For pA-DamID on the knock-out clones and clone 5, the scores were calculated in a window 10kb up and downstream from the IPR. Except for the different window size, pA-DamID scores were calculated similar to the overall DamID scores. Z-scores for pA-DamID were calculated using the mean and standard deviation of the pA-DamID score for 20kb binned tracks over the whole genome using the same formula as for the individual IPR’s.

The following formula was used to calculate Z-scores:

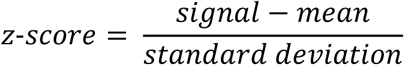

### Chromatin immunoprecipitation analysis of IPRs

Chromatin immunoprecipitation was performed as described (Schmidt et al., 2009). Main steps and modifications are described here. 50 µl protein A DynabeadsTM were precleared with 0.5% BSA, 5 µl of specific antibody (H3K4me1: Abcam ab8895; H3K27me3: Active Motif 39155; H3K27ac: Active Motif 39133) and beads incubated at 4°C overnight. 10 million clone 5 cells were fixed at a final concentration of 1% formaldehyde for 10 minutes. Fixation was quenched with 125 mM glycine for 5 minutes and a PBS wash. After nuclear extraction, chromatin was sonicated (∼8 cycles 30 sec on / 30 sec off in BioRuptor Pico), Triton-X-100 added to a final concentration of 1% and centrifuged to remove cell debris. Antibody coupled beads were washed with 0.5% BSA in PBS, chromatin was added (5% was kept as input) and rotated overnight at 4°C. Beads were washed 10 times with RIPA buffer and once with TBS. After last wash, 200 µl of elution buffer was added and samples eluted and de-crosslinked at 65°C overnight. 200 µl of TE buffer and 0.9 µl of 10mg/ml RNAse A was added the samples and were incubated at 37°C for 1 hour and with 4 µl of 20mg/ml Proteinase K at 55°C for 2 hours. DNA was extracted by phenol:chloroform extraction and resuspended in 50 µl of 10nM Tris-HCl. IPR barcodes were collectively amplified using primers the two step PCR described above with slight modifications. For indelPCR1, 100ng DNA was taken from input samples and same input volume added from pull-downs. IndelPCR1 was performed in a final volume of 50 µl, with 25µl MyTaq HS Red mix and 0.5 µM of each primer (TAC0007 and TAC0012). 5µl of indelPCR1 was taken as input for indelPCR2. This PCR was performed in a final volume of 50µl, with 25µl of MyTaq Red mix and 0.5 µM of each primer (TAC0009 and TAC0159) for 12 PCR cycles (3 cycles with 58°C annealing followed by 9 cycles with 65°C annealing). PCR products were pooled, purified as described above, and sequenced on an Illumina MiSeq.

### Time series analyses

Indels in clone 5 were identified and counted as described above. Indel frequencies and MMEJ:NHEJ balance was calculated before averaging across replicates. Only IPRs mapped to a single genomic location were used (19 total). Sigmoid curves were fitted to time series data using the following formula:

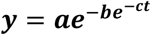

Where t is time and a, b and c are parameters that determine the shape and plateau of the curve. For the decay of wild-type sequence over time, the ratio was fitted as 1-y. Fitting was done using the *nls* package in R. Starting values 20, 10 and 0.1 were used for fitting of the parameters a, b and c, respectively.

### Data availability

Processed data is available at https://osf.io/cywxd/.

**Supplementary Table 1.** Related to Methods. Epigenome ChIP datasets used in this study.

| chip_source | chip_target | chip_id | sra_chip | sra_input | PMID |
| --- | --- | --- | --- | --- | --- |
| **Bernstein2011** | H2AFZ | GSM733786 | SRR227661,<br>SRR227662 | SRR227650,<br>SRR5331211,<br>SRR5331212,<br>SRR5331213 | 22955616 |
| **Bernstein2012** | EZH2 | GSM1003576 | SRR568431,<br>SRR568432 | SRR227650,<br>SRR5331211,<br>SRR5331212,<br>SRR5331213 | 22955616 |
| **Bernstein2012** | H3K79me2 | GSM733653 | SRR227378,<br>SRR227379 | SRR227650,<br>SRR5331211,<br>SRR5331212,<br>SRR5331213 | 22955616 |
| **Minoda2018** | H4K5acK8ac | GSE113635 | SRR7070730,<br>SRR7070731 | SRR7070732 | 30503705 |
| **Salzberg2017** | H3K9me2 | GSM1846169,<br>GSM2152591 | SRR2148301,<br>SRR3503783 | SRR2148307 | 28301528 |
| **Schmidl2015** | CTCF | GSM1782717,<br>GSM1782718 | SRR2085871,<br>SRR2085872 | SRR2085882,<br>SRR2085883,<br>SRR2085884,<br>SRR2085885,<br>SRR2085886 | 26280331 |
| **Schmidl2015** | H3K27ac | GSM1782721,<br>GSM1782722 | SRR2085875,<br>SRR2085876 | SRR2085882,<br>SRR2085883,<br>SRR2085884,<br>SRR2085885,<br>SRR2085886 | 26280331 |
| **Schmidl2015** | H3K27me3 | GSM1782749,<br>GSM1782750 | SRR2085903,<br>SRR2085904 | SRR2085882,<br>SRR2085883,<br>SRR2085884,<br>SRR2085885,<br>SRR2085886 | 26280331 |
| **Schmidl2015** | H3K36me3 | GSM1782723,<br>GSM1782724 | SRR2085877,<br>SRR2085878 | SRR2085882,<br>SRR2085883,<br>SRR2085884,<br>SRR2085885,<br>SRR2085886 | 26280331 |
| **Shah2018** | H3K4me1 | GSM2773392 | SRR6010166 | SRR6010181 | 30244833 |
| **Shah2018** | H3K4me1 | GSM2773394 | SRR6010168 | SRR6010181 | 30244833 |
| **Shah2018** | H3K4me1 | GSM2773396 | SRR6010170 | SRR6010181 | 30244833 |
| **Shah2018** | H3K4me2 | GSM2773399 | SRR6010173 | SRR6010181 | 30244833 |
| **Shah2018** | H3K4me2 | GSM2773400 | SRR6010174 | SRR6010181 | 30244833 |
| **Shah2018** | H3K4me3 | GSM2773401 | SRR6010175 | SRR6010181 | 30244833 |
| **Shah2018** | H3K4me3 | GSM2773403 | SRR6010177 | SRR6010181 | 30244833 |
| **Shah2018** | H3K4me3 | GSM2773404 | SRR6010178 | SRR6010181 | 30244833 |
| **Shah2018** | H3K4me3 | GSM2773406 | SRR6010180 | SRR6010181 | 30244833 |
| **Snyder2012** | POL2AS2 | GSM935402 | SRR502194,<br>SRR502195 | SRR502641 | 22955616 |
| **Snyder2012** | SMC3 | GSM935310 | SRR502001,<br>SRR502002 | SRR502641 | 22955616 |
| **Snyder2016** | POL2 | GSE91721 | SRR5111542,<br>SRR5111543 | SRR5111209,<br>SRR5111210 | 22955616 |
| **Snyder2017** | HDAC1 | GSE105837 | SRR6213961,<br>SRR6213962 | SRR5111209,<br>SRR5111210 | 22955616 |
| **Snyder2017** | HDAC2 | GSE91451 | SRR5111049,<br>SRR5111050 | SRR5111209,<br>SRR5111210 | 22955616 |
| **Snyder2018** | HDAC3 | GSE127356 | SRR8659957,<br>SRR8659958 | SRR5111896,<br>SRR5111897 | 22955616 |

**Supplementary Table 2.**
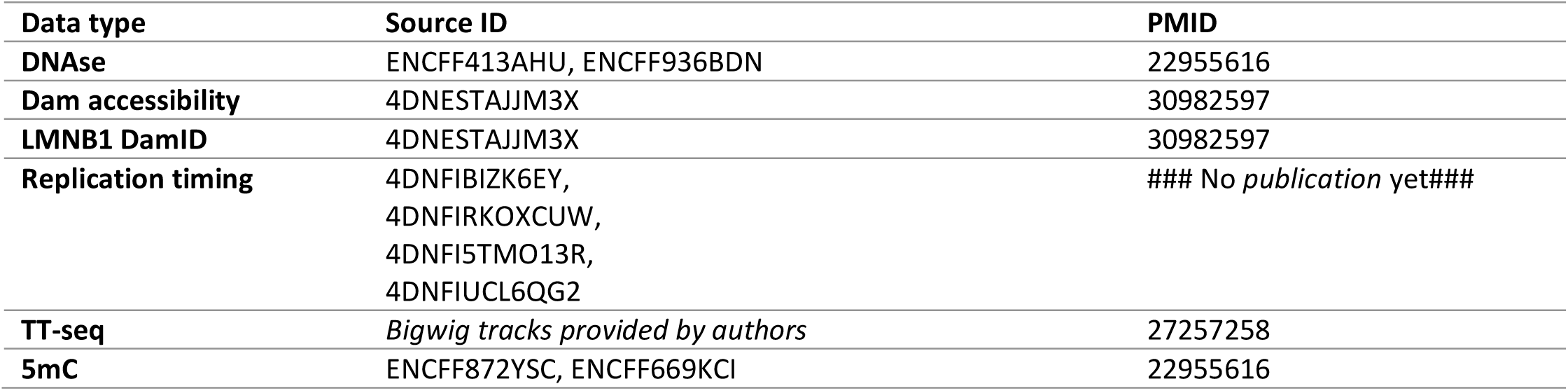
Related to Methods.

**Supplementary Table 3.**
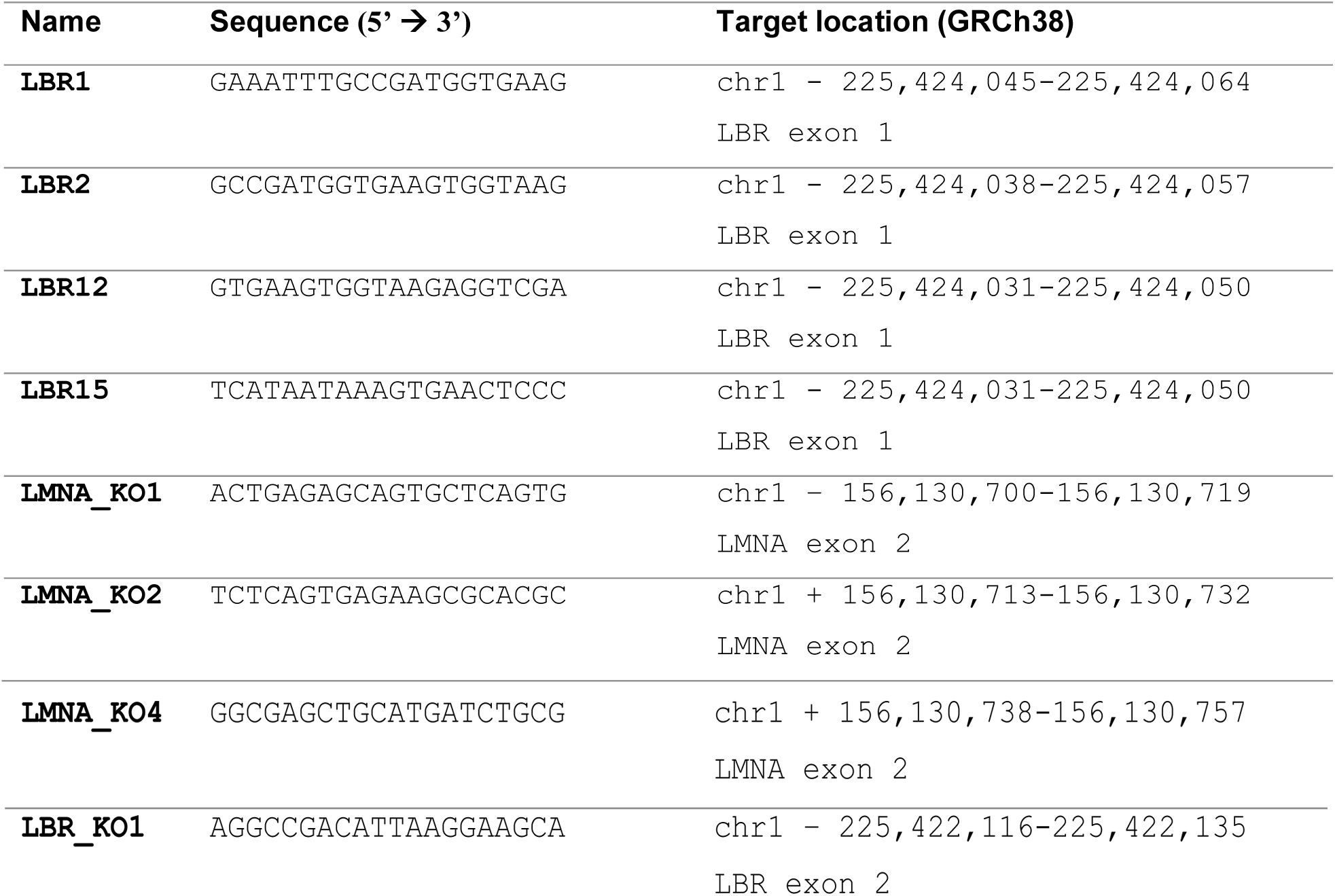
Related to Methods.

**Supplementary Table 4.**
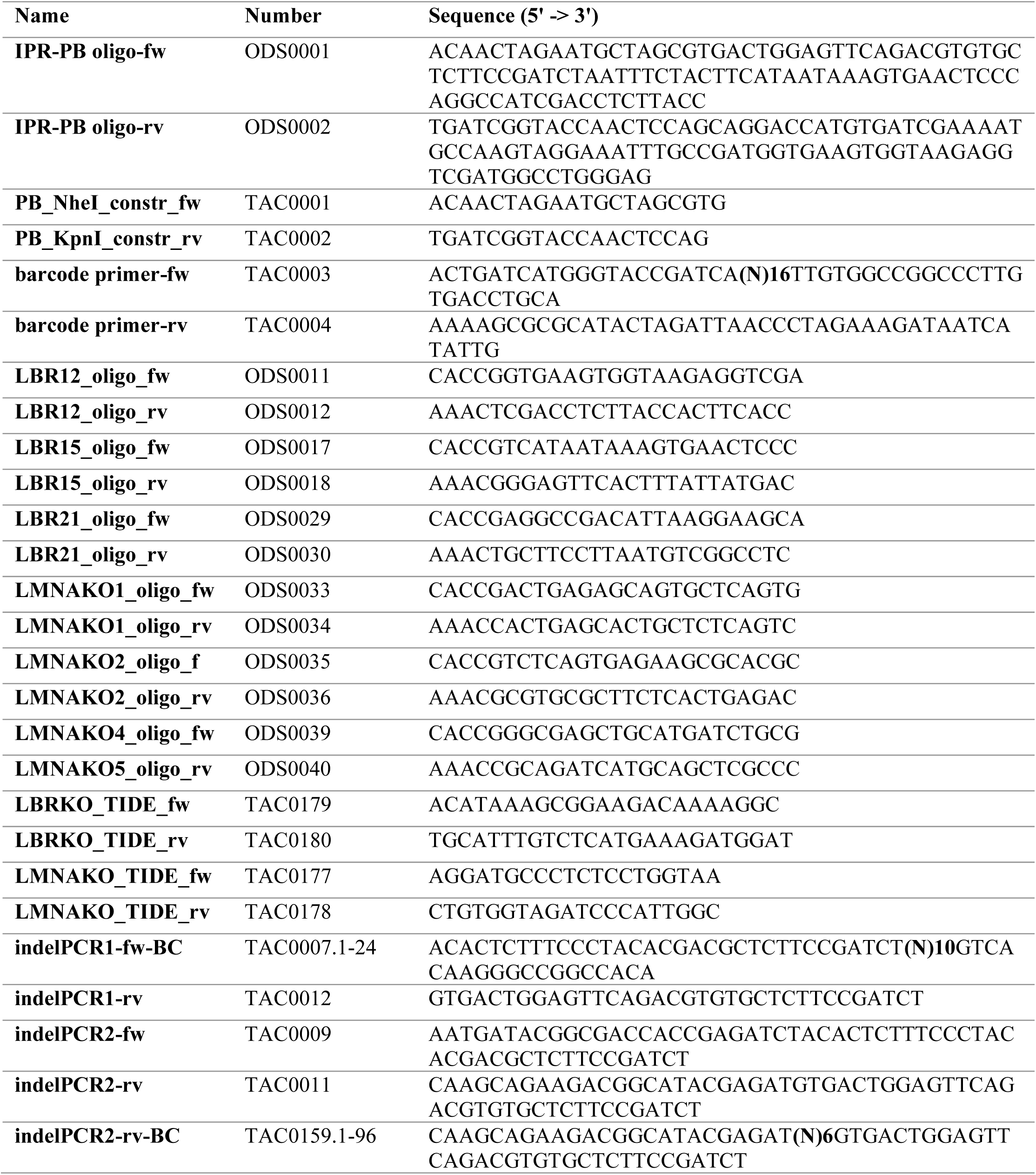
Related to Methods.

## SUPPLEMENTARY FIGURE LEGENDS

**Supplementary Figure S1:**
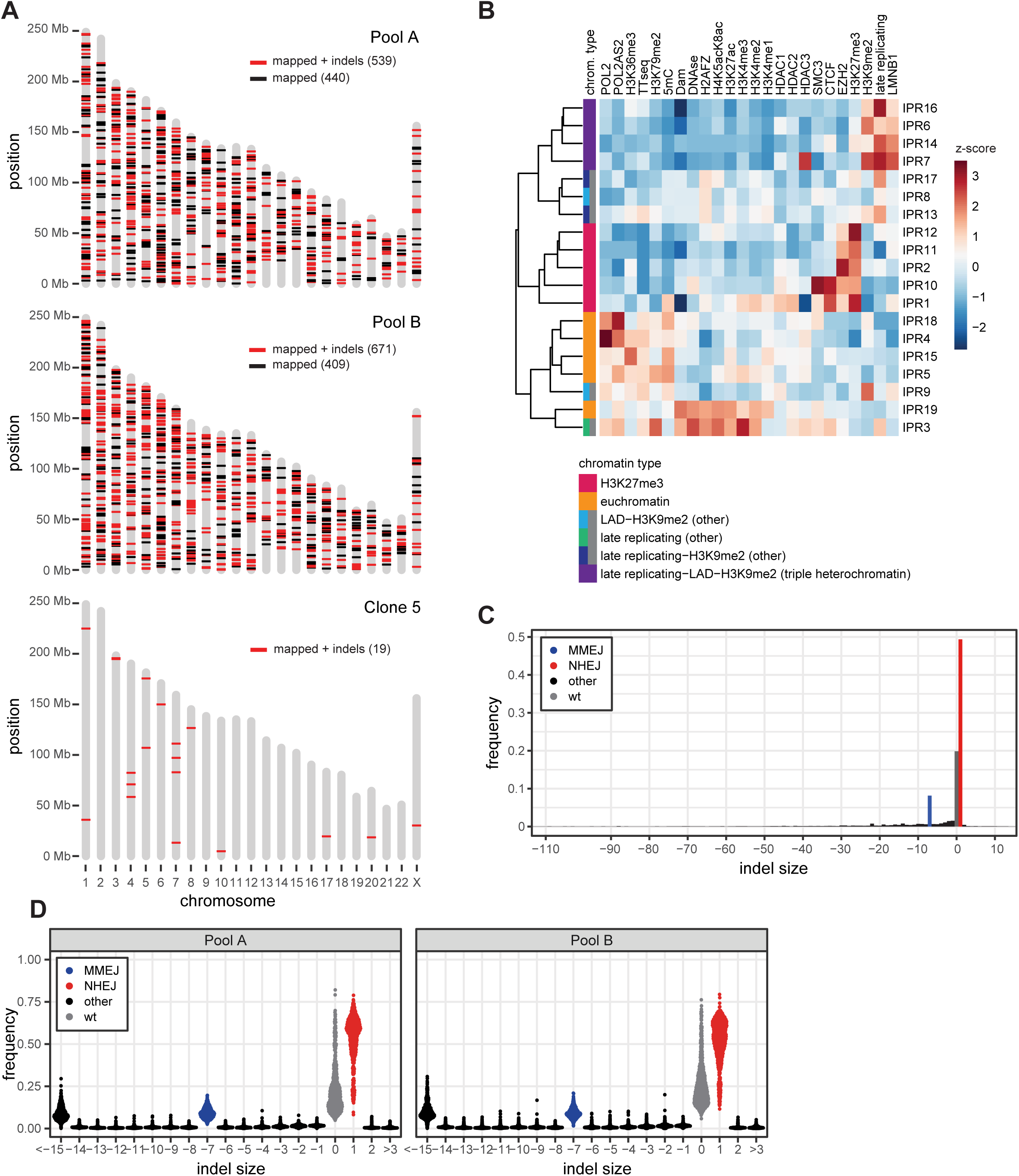
genomic location, chromatin context and indel frequencies of IPRs. (**A**) Genomic locations of the mapped IPRs in each of the two cell pools (Pools A and B) and in clone 5. Red: uniquely mapped IPRs with indel data that passed the quality criteria (also shown in **Figure 2A**); black: uniquely mapped IPRs that did not pass the indel quality criteria. (see Methods) (**B**) Heatmap of chromatin features at each IPR integration site in clone 5. Levels of chromatin features are represented as z-scores (see Methods). IPRs are clustered based on similarities of the z-scores. (**C**) Median indel frequencies across all IPRs; same data as Figure 2B but plotted over a wider range of indel sizes to illustrate that large indels are rare compared to −7 and +1. (**D**) Frequencies of all indel sizes as in **Figure 2C** but split on each pool. Data for 539 (pool A) and 690 (pool B) IPRs. Data are average of 2-6 independent replicates.

**Supplementary Figure S2.**
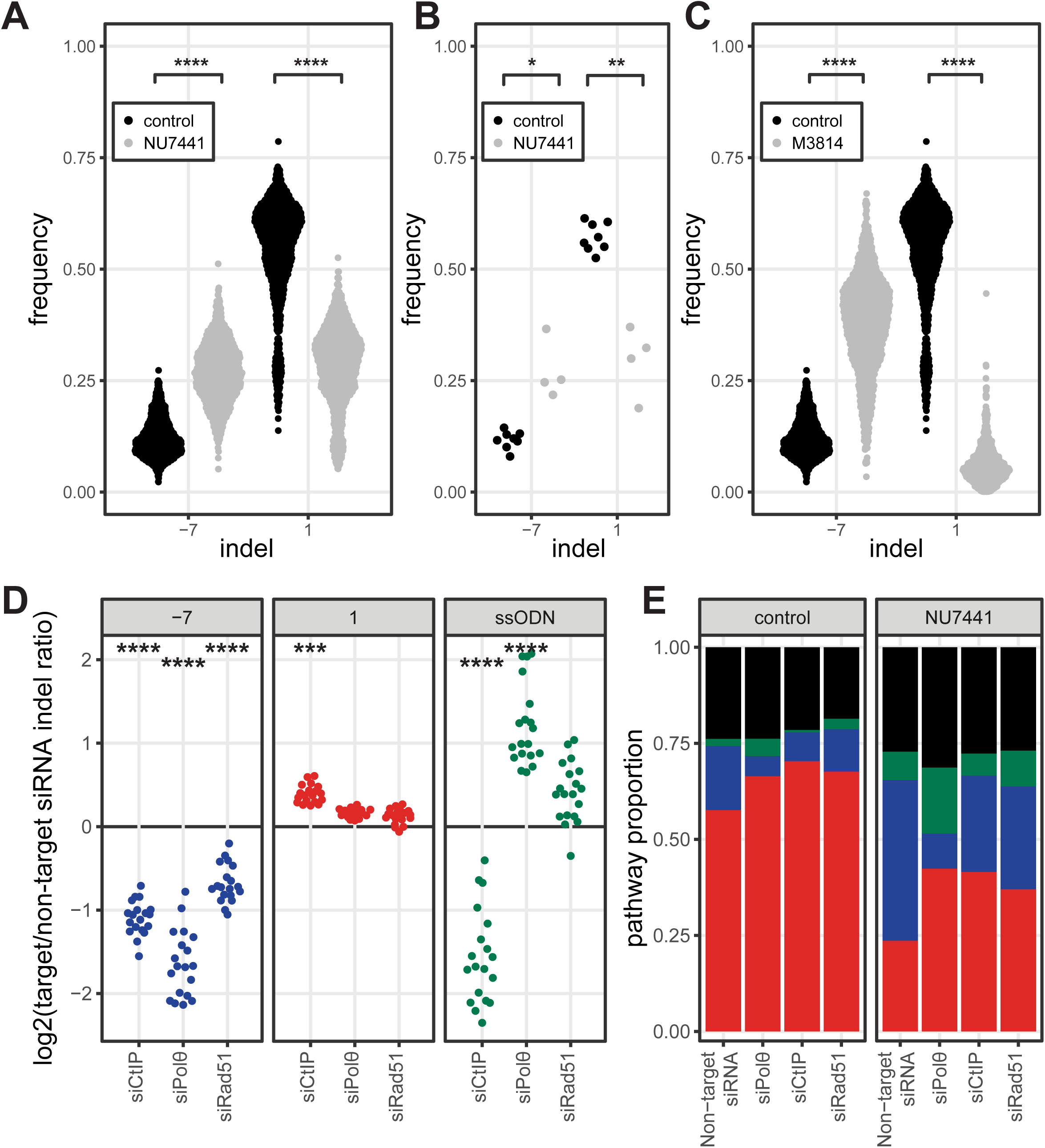
Characterization of pathways that generate reporter indels. **(A-C)** Effects of chemical inhibition of DNAPKcs on indel frequencies. (**A**) Frequency of +1 and −7 indels for all IPRs in the cell pools treated with the DNAPKcs inhibitor NU7441 (gray, mean of n = 5) or with DMSO control (black, mean of n = 6). (**B**) Same data as in A, but now shown as median indel frequencies of all IPRs, for each replicate experiment. (**C**) Frequency of +1 and −7 indels for all IPRs in the cell pools treated with the DNAPKcs inhibitor M3814 (gray, mean of n = 2) or with DMSO control (black, n = 2). **(D-E)** Effect of knockdown of various DSB repair proteins on indel frequencies. (**D**) Log_2_ fold-change in the frequencies of the −7, +1 and ssODN-induced +2 indels at all 19 IPRs in clone 5, after siRNA-mediated knockdown of the indicated proteins compared to a control siRNA (n = 2). (**E**) Relative activity of each pathway for all 19 barcodes in clone 5 combined, after indicated siRNA treatments. Red: +1 insertion (NHEJ); blue: −7 deletion (MMEJ); green: +2 insertion due to SSTR; black: other indels. Right-hand panel shows data from cells treated with NU7441 (1 µM), left-hand panel shows data from control cells. Asterisks in panels **A-D** denote adjusted p-values: * p < 0.05; ** p < 0.01; *** p < 0.001; **** p < 0.0001.

**Supplementary Figure S3:**
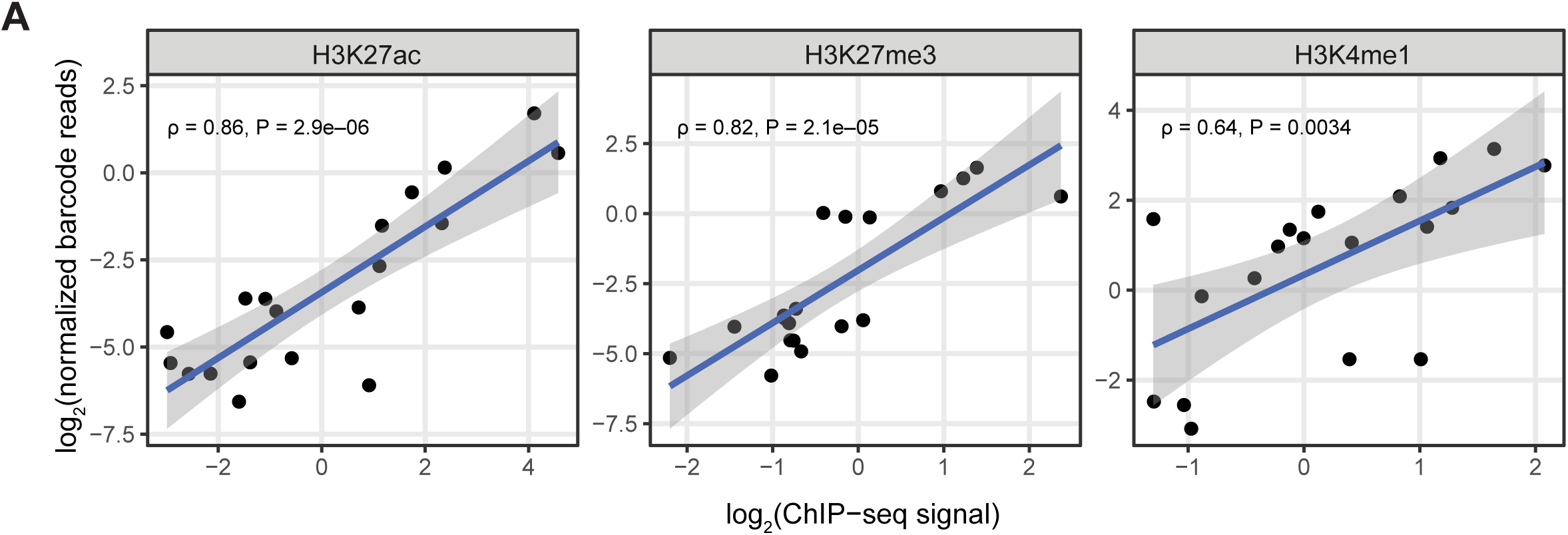
IPRs generally adopt the chromatin state of their integration site. **(A**) Horizontal axes: log_2_ normalized signal of the indicated histone modifications in a window of 2kb centered on the IPR integration site (n = 2), according to public ChIP-seq data from K562 cells (see **Table S1**). Vertical axes: log_2_ normalized barcode reads of 19 IPRs in clone 5 after ChIP of the indicated histone modifications, followed by PCR amplification of the barcodes and Illumina sequencing. Data are average of two independent replicates. Solid lines and shading show linear regression fits with 95% confidence intervals. Rho is Spearman’s rank correlation coefficient.

**Supplementary Figure S4:**
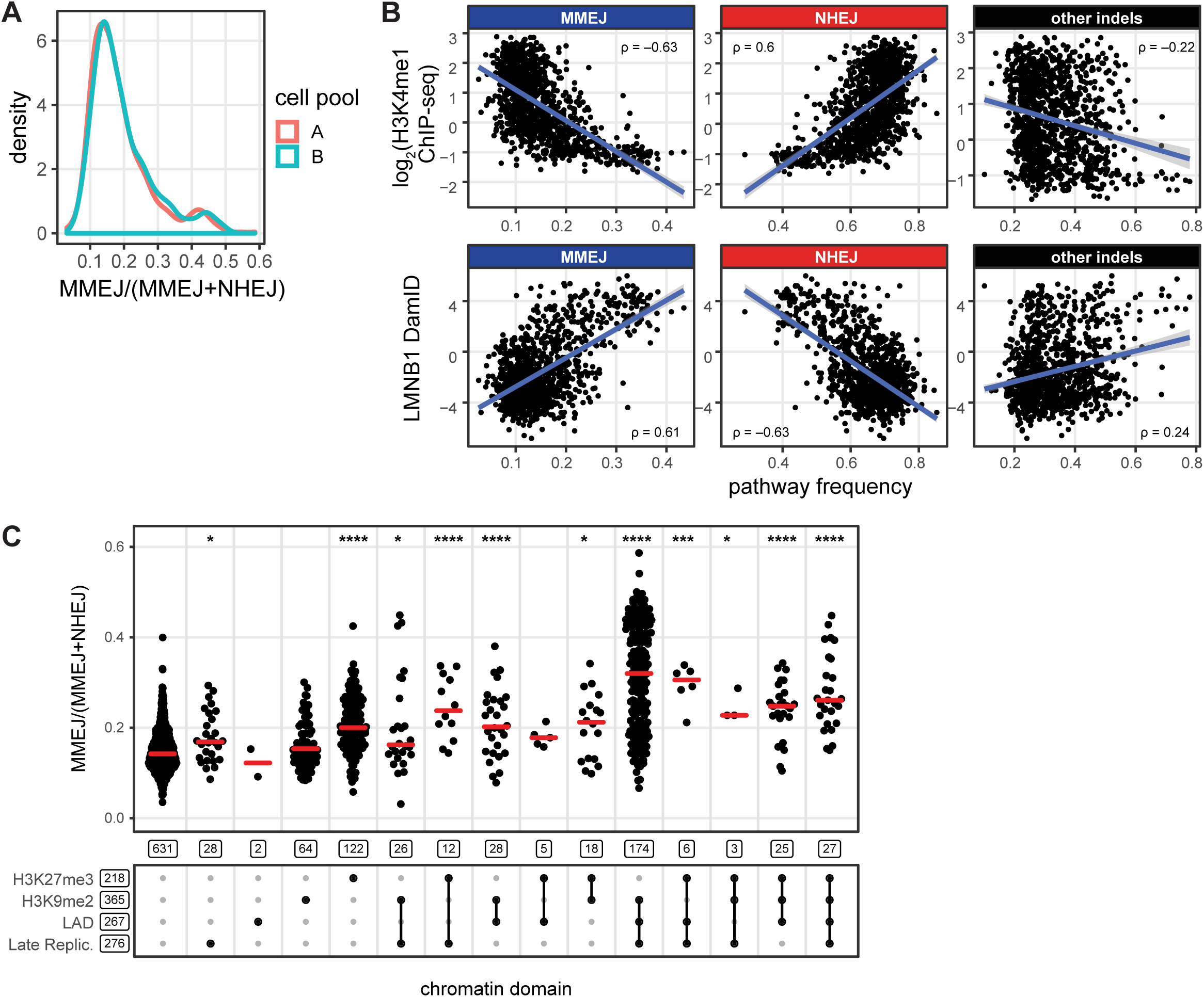
Correlations between pathway usage and chromatin features. (**A**) Scatterplots of the relative indel frequencies in the IPR cell pools versus local log_2_ H3K4me1 ChIP-seq signal (top row) or log_2_ Lamin B1 DamID signal (bottom row) at the IPR integration sites. Rho is Pearson’s correlation coefficient; solid lines and shading show linear regression fits with 95% confidence intervals. (**B**) Density plots of the MMEJ:NHEJ balance for each pool separately. (**B**) Same as **Figure 4D**, but now including heterochromatin types with 20 or fewer IPRs.

**Supplementary Figure S5:**
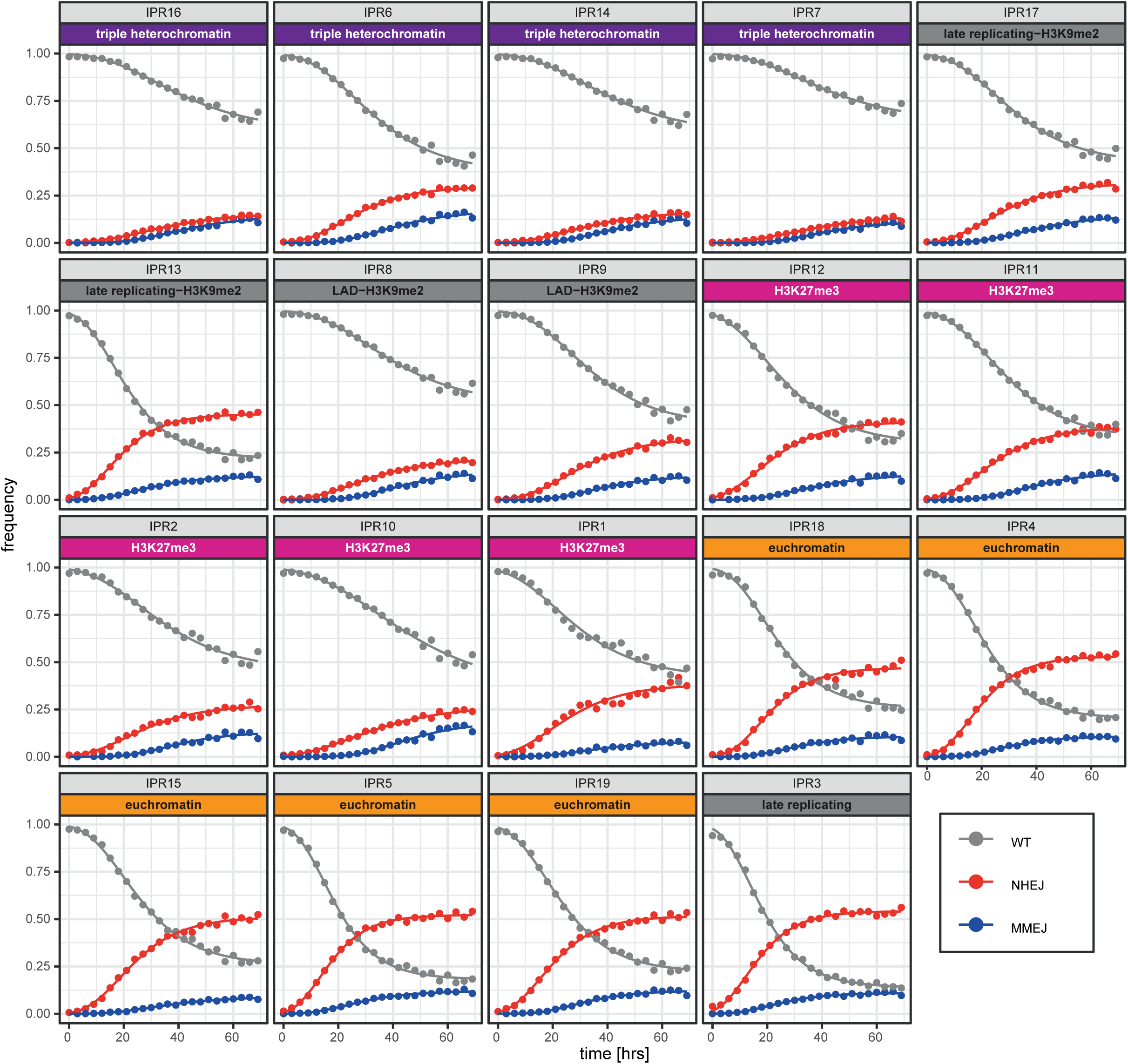
Time series of −7 and +1 indel accumulation for all IPRs in clone 5. Time curves of the +1 insertion (red) and −7 deletion (blue) for all 19 individual IPRs. See legend Figure 5A. Triple heterochromatin is the combination of late replicating, LAD and H3K9me2.

**Supplementary Figure S6:**
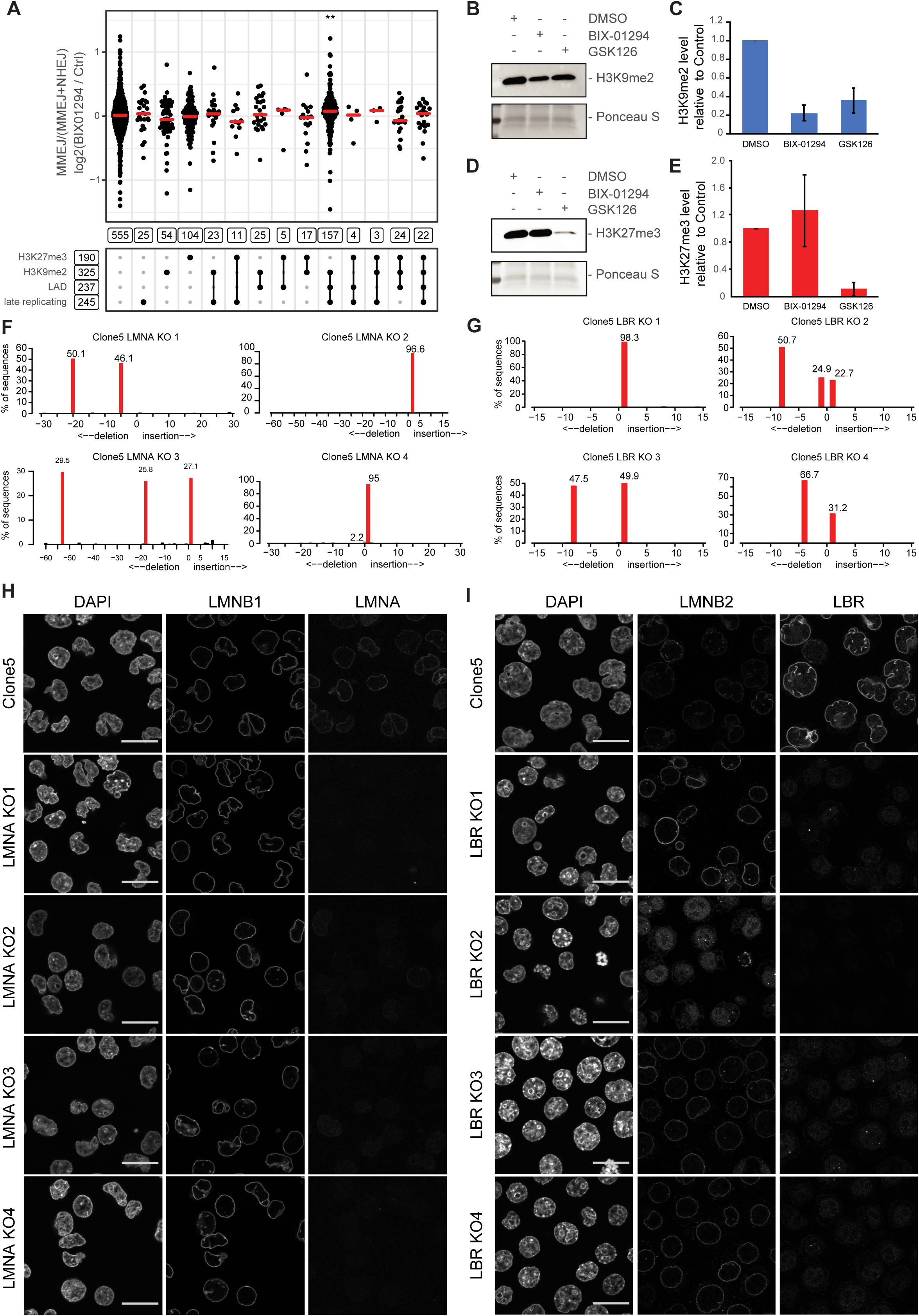
Inhibitor and knockdown validations. (**A**) Log_2_ fold-change of MMEJ:NHEJ balance in BIX01294 treated cells (n = 2) compared to non-treated cells for 1029 IPRs divided by heterochromatin domains (n = 2-8). Wilcoxon test compared to non-heterochromatin IPRs (most left column), * p < 0.05, ** p < 0.01, *** p < 0.001, **** p < 0.0001. (**B**) Western blot of H3K9me2 in clone 5 cells after treatment with either 1 µM GSK126 or 500 nM BIX01294. (**C**) Quantification of Western blots (mean of two independent replicates, error bars show S.D.), normalized to H3K9me2 levels in control cells and red ponceau staining for protein content. (**D-E**) Same as B and C but for H3K27me3. (**F-G**) Indel patterns inside the *LMNA* (E) and *LBR* (F) genes in respective knockout sub-clones that were derived from clone 5, showing frameshifts (i.e., indel sizes that are not multiples of three) in all alleles. Note that chromosomes in K562 cells can be tri- or tetraploid. Indel spectra were obtained by TIDE (Brinkman et al., 2014). (**H**) DAPI and immunostaining of LMNB1 and LMNA in wild-type clone 5 cells and in the four independent LMNA knockout cell lines. (**I**) Immunostaining of LMNB1 (red) and LBR (red) in wild-type clone 5 (WT) cells and in the four independent LBR knockout cell lines. Scale bar in G-H is 20 μm.

**Supplementary Figure S7:**
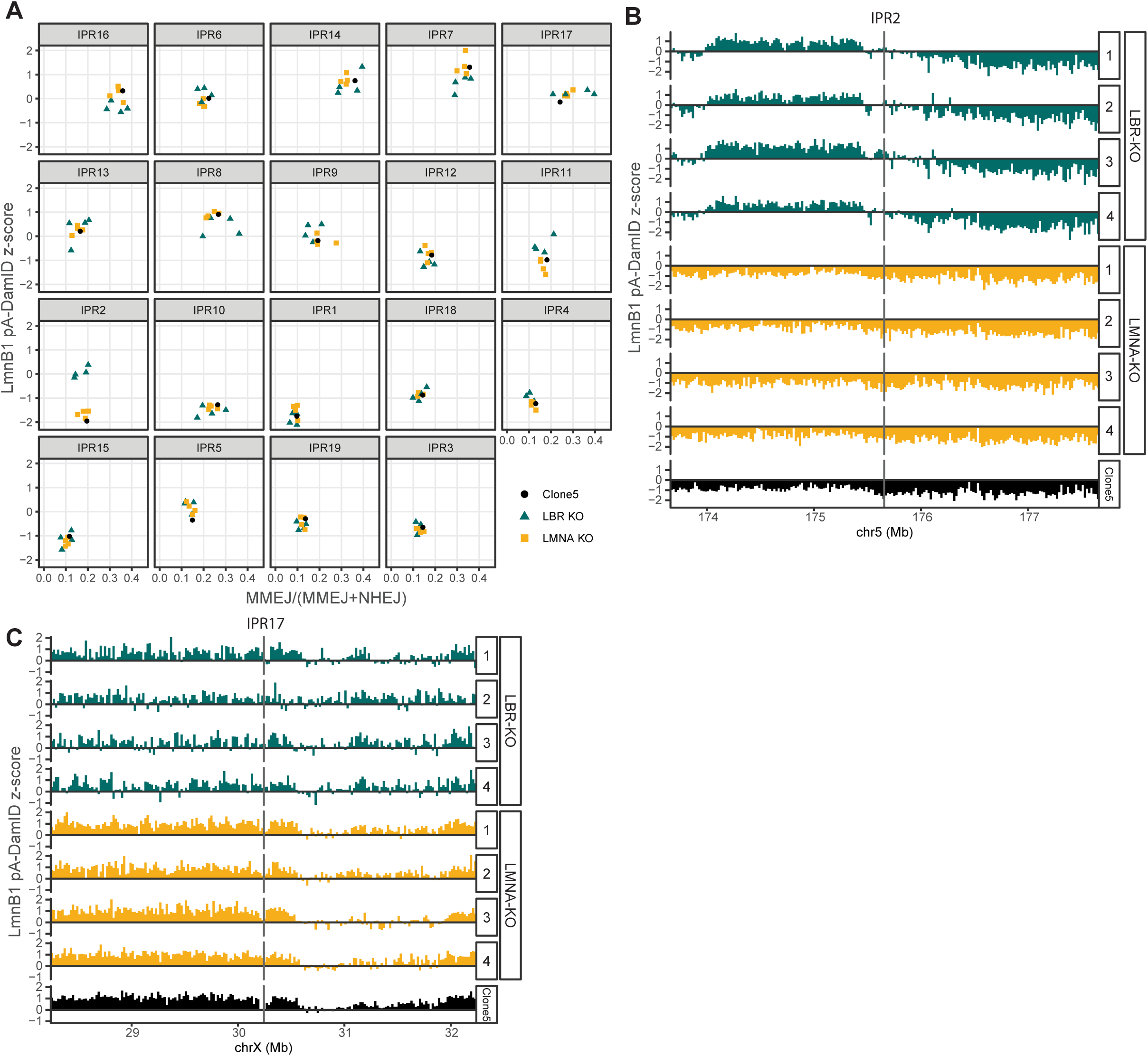
LMNA and LBR knockout in clone 5. (**D**) Scatterplot of MMEJ:NHEJ balance compared to LMNB1 pA-DamID z-score for each IPR, 10kb up and downstream of the IPR. Black circle is clone5, green triangles are the LBR KO clones and yellow squares are LMNA KO clones. (**A**) Z-score of pA-DamID tracks for LMNB1 centered on IPR17 with 2Mb up and downstream. (n = 2) (**C**) Same as in B but for IPR2.

**Supplementary Figure S8:**
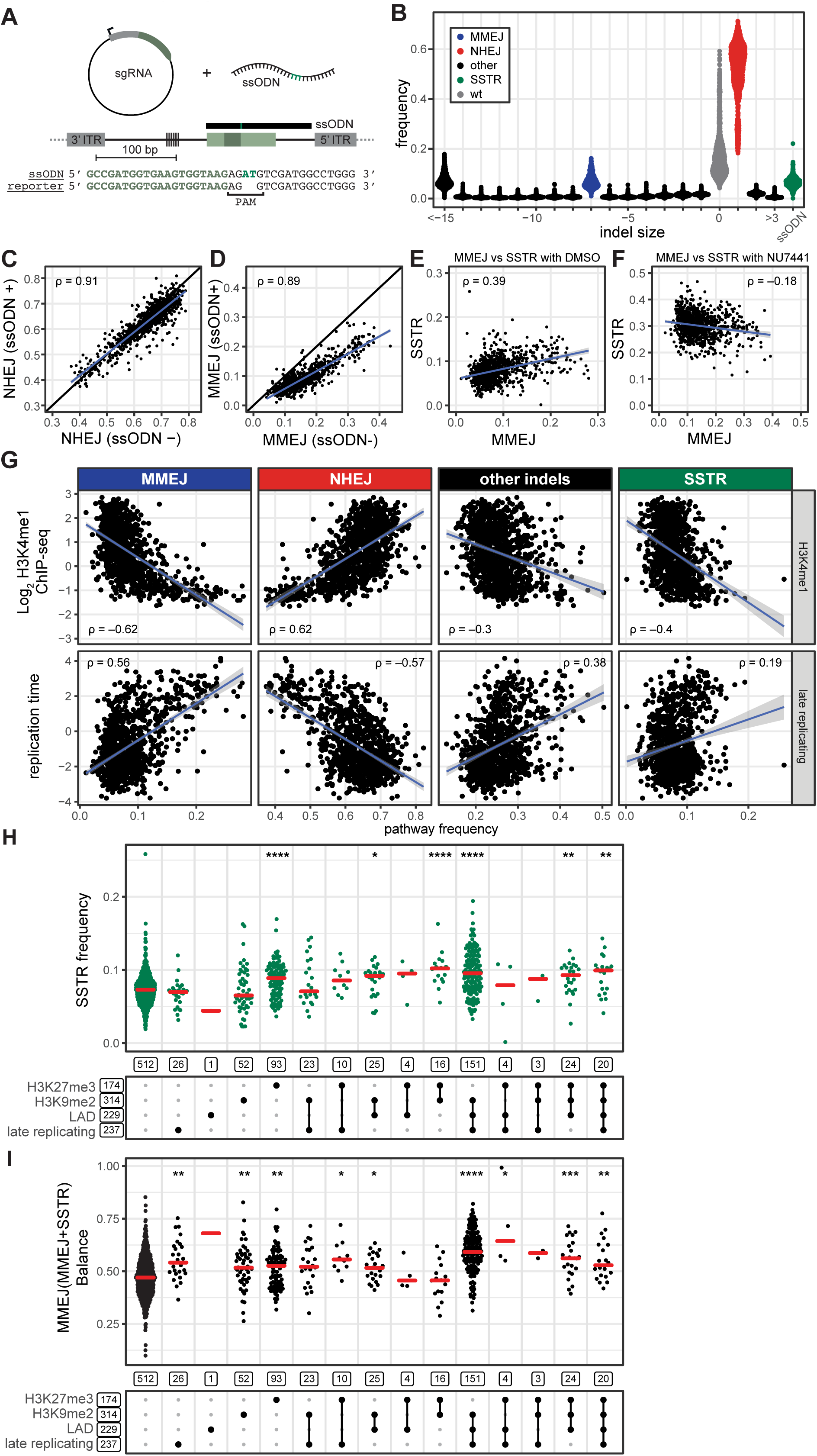
Chromatin context effects on the SSTR pathway. **(A)** Schematic of the strategy to probe SSTR simultaneously with NHEJ and MMEJ. Prior to Cas9 activation, the ssODN is co-transfected with a plasmid that encodes the LBR2 sgRNA. The ssODN (black bar) covers the reporter sequence but not the barcode, and contains a 2 bp insertion (green) at the PAM. **(B)** Indel frequencies generated by NHEJ, MMEJ and SSTR in 965 IPRs in the two cell pools, 64 hours after Cas9 activation (average of two replicate experiments). N = 2-3 (**C-D**) Comparison of +1 (NHEJ; panel **C**) and −7 (MMEJ, panel **D**) indel frequencies in all IPRs in cell pools in the presence (+) or absence (-) of the ssODN. Black line: diagonal. (**E**) Comparison of MMEJ and SSTR frequencies across all IPRs in cell pools treated with the ssODN. (**F**) Same as (**E**) but in the presence of NU7441. (**G-H**) Scatterplots of the relative indel frequencies (proportion of all indels) versus (**G**) log_2_ H3K4me1 ChIP-seq signal and (**H**) log_2_ replication timing (positive values: late, negative values: early). *R* in panels C-G are Pearson’s correlation coefficients; blue lines with shading are linear regression fits with 95% confidence interval. (**I-J**) Same as Figures 7C-D, but now including heterochromatin types with 20 or fewer IPRs.

**Supplementary Figure S9:**
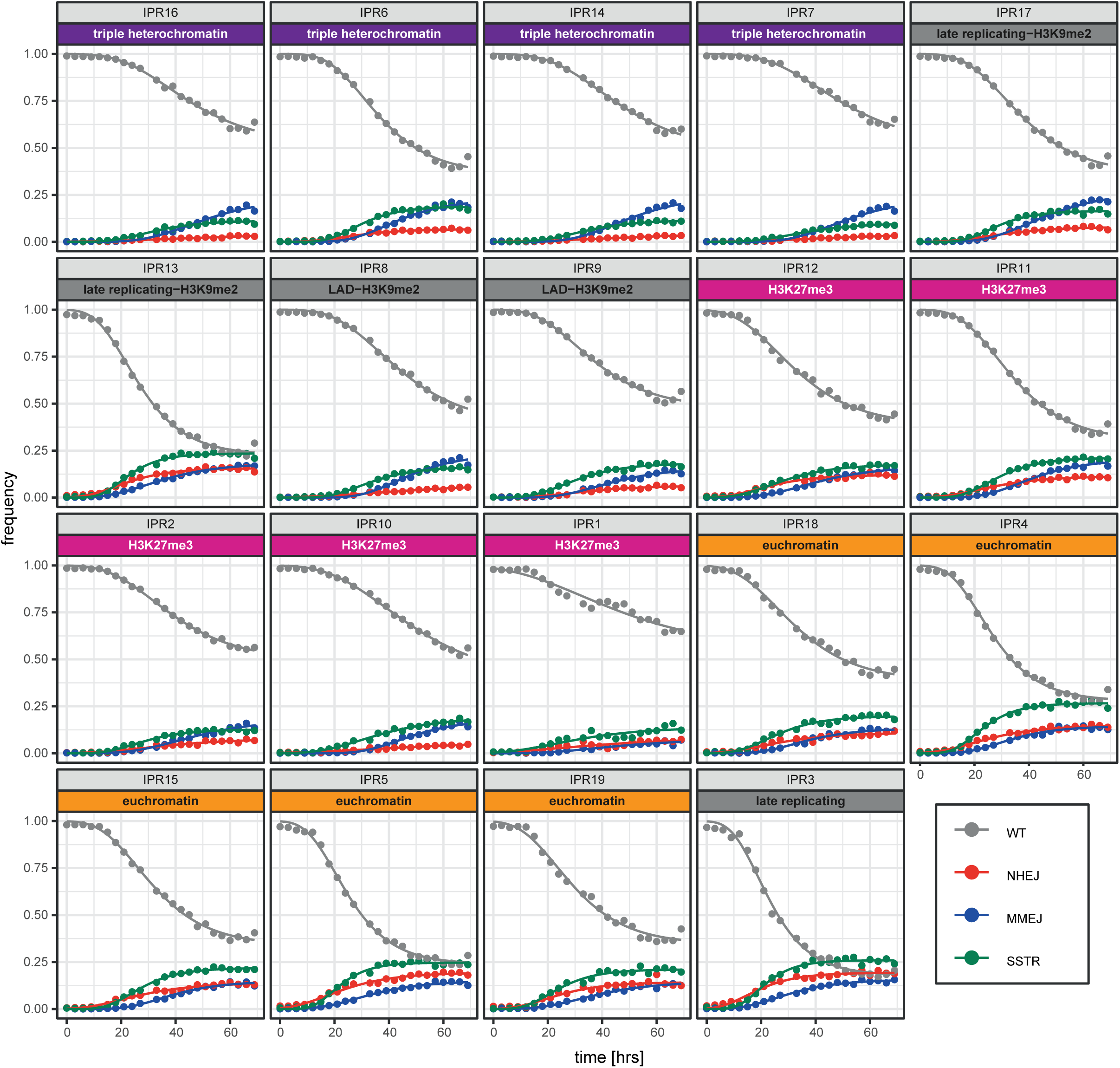
Time series of indel accumulation, including SSTR. Time curves as in **Figure 7E**, but now for all 19 individual IPRs in clone 5 in the presence of ssODN to probe SSTR. Cells were treated with NU7441 to reduce the contribution of NHEJ. See legend **Figure 7E** for details. Triple heterochromatin is the combination of late replicating, LAD and H3K9me2.

## REFERENCES

Akhtar, W., de Jong, J., Pindyurin, A.V., Pagie, L., Meuleman, W., de Ridder, J., Berns, A., Wessels, L.F., van Lohuizen, M., and van Steensel, B. (2013). Chromatin position effects assayed by thousands of reporters integrated in parallel. Cell 154, 914–927.

Akhtar, W., Pindyurin, A.V., de Jong, J., Pagie, L., Ten Hoeve, J., Berns, A., Wessels, L.F., van Steensel, B., and van Lohuizen, M. (2014). Using TRIP for genome-wide position effect analysis in cultured cells. Nat Protoc 9, 1255–1281.

Alagoz, M., Katsuki, Y., Ogiwara, H., Ogi, T., Shibata, A., Kakarougkas, A., and Jeggo, P. (2015). SETDB1, HP1 and SUV39 promote repositioning of 53BP1 to extend resection during homologous recombination in G2 cells. Nucleic Acids Res 43, 7931–7944.

Allen, F., Crepaldi, L., Alsinet, C., Strong, A.J., Kleshchevnikov, V., De Angeli, P., Palenikova, P., Khodak, A., Kiselev, V., Kosicki, M., et al. (2018). Predicting the mutations generated by repair of Cas9-induced double-strand breaks. Nat Biotechnol.

Aymard, F., Bugler, B., Schmidt, C.K., Guillou, E., Caron, P., Briois, S., Iacovoni, J.S., Daburon, V., Miller, K.M., Jackson, S.P., et al. (2014). Transcriptionally active chromatin recruits homologous recombination at DNA double-strand breaks. Nat Struct Mol Biol 21, 366–374.

Baldeyron, C., Soria, G., Roche, D., Cook, A.J., and Almouzni, G. (2011). HP1alpha recruitment to DNA damage by p150CAF-1 promotes homologous recombination repair. J Cell Biol 193, 81–95.

Banaszynski, L.A., Chen, L.C., Maynard-Smith, L.A., Ooi, A.G., and Wandless, T.J. (2006). A rapid, reversible, and tunable method to regulate protein function in living cells using synthetic small molecules. Cell 126, 995–1004.

Brinkman, E.K., Chen, T., Amendola, M., and van Steensel, B. (2014). Easy quantitative assessment of genome editing by sequence trace decomposition. Nucleic Acids Res 42, e168.

Brinkman, E.K., Chen, T., de Haas, M., Holland, H.A., Akhtar, W., and van Steensel, B. (2018). Kinetics and Fidelity of the Repair of Cas9-Induced Double-Strand DNA Breaks. Mol Cell 70, 801–813 e806.

Carvalho, S., Vitor, A.C., Sridhara, S.C., Martins, F.B., Raposo, A.C., Desterro, J.M., Ferreira, J., and de Almeida, S.F. (2014). SETD2 is required for DNA double-strand break repair and activation of the p53-mediated checkpoint. Elife 3, e02482.

Chakrabarti, A.M., Henser-Brownhill, T., Monserrat, J., Poetsch, A.R., Luscombe, N.M., and Scaffidi, P. (2019). Target-Specific Precision of CRISPR-Mediated Genome Editing. Mol Cell 73, 699–713 e696.

Chan, S.H., Yu, A.M., and McVey, M. (2010). Dual roles for DNA polymerase theta in alternative end-joining repair of double-strand breaks in Drosophila. PLoS Genet 6, e1001005.

Chang, H.H.Y., Pannunzio, N.R., Adachi, N., and Lieber, M.R. (2017). Non-homologous DNA end joining and alternative pathways to double-strand break repair. Nat Rev Mol Cell Biol 18, 495–506.

Chapman, J.R., Taylor, M.R., and Boulton, S.J. (2012). Playing the end game: DNA double-strand break repair pathway choice. Mol Cell 47, 497–510.

Chen, W., McKenna, A., Schreiber, J., Haeussler, M., Yin, Y., Agarwal, V., Noble, W.S., and Shendure, J. (2019). Massively parallel profiling and predictive modeling of the outcomes of CRISPR/Cas9-mediated double-strand break repair. Nucleic Acids Res 47, 7989–8003.

Chen, X., Rinsma, M., Janssen, J.M., Liu, J., Maggio, I., and Goncalves, M.A. (2016). Probing the impact of chromatin conformation on genome editing tools. Nucleic Acids Res 44, 6482–6492.

Chen, Y., Zhang, Y., Wang, Y., Zhang, L., Brinkman, E.K., Adam, S.A., Goldman, R., van Steensel, B., Ma, J., and Belmont, A.S. (2018). Mapping 3D genome organization relative to nuclear compartments using TSA-Seq as a cytological ruler. J Cell Biol 217, 4025–4048.

Chiolo, I., Minoda, A., Colmenares, S.U., Polyzos, A., Costes, S.V., and Karpen, G.H. (2011). Double-strand breaks in heterochromatin move outside of a dynamic HP1a domain to complete recombinational repair. Cell 144, 732–744.

Clouaire, T., and Legube, G. (2015). DNA double strand break repair pathway choice: a chromatin based decision? Nucleus 6, 107–113.

Clouaire, T., and Legube, G. (2019). A Snapshot on the Cis Chromatin Response to DNA Double-Strand Breaks. Trends Genet 35, 330–345.

Clouaire, T., Rocher, V., Lashgari, A., Arnould, C., Aguirrebengoa, M., Biernacka, A., Skrzypczak, M., Aymard, F., Fongang, B., Dojer, N., et al. (2018). Comprehensive Mapping of Histone Modifications at DNA Double-Strand Breaks Deciphers Repair Pathway Chromatin Signatures. Mol Cell 72, 250–262 e256.

Clowney, E.J., LeGros, M.A., Mosley, C.P., Clowney, F.G., Markenskoff-Papadimitriou, E.C., Myllys, M., Barnea, G., Larabell, C.A., and Lomvardas, S. (2012). Nuclear aggregation of olfactory receptor genes governs their monogenic expression. Cell 151, 724–737.

Corrales, M., Rosado, A., Cortini, R., van Arensbergen, J., van Steensel, B., and Filion, G.J. (2017). Clustering of Drosophila housekeeping promoters facilitates their expression. Genome Res 27, 1153–1161.

Daer, R.M., Cutts, J.P., Brafman, D.A., and Haynes, K.A. (2017). The Impact of Chromatin Dynamics on Cas9-Mediated Genome Editing in Human Cells. ACS Synth Biol 6, 428–438.

Daugaard, M., Baude, A., Fugger, K., Povlsen, L.K., Beck, H., Sorensen, C.S., Petersen, N.H., Sorensen, P.H., Lukas, C., Bartek, J., et al. (2012). LEDGF (p75) promotes DNA-end resection and homologous recombination. Nat Struct Mol Biol 19, 803–810.

Deng, S.K., Gibb, B., de Almeida, M.J., Greene, E.C., and Symington, L.S. (2014). RPA antagonizes microhomology-mediated repair of DNA double-strand breaks. Nat Struct Mol Biol 21, 405–412.

DeWitt, M.A., Magis, W., Bray, N.L., Wang, T., Berman, J.R., Urbinati, F., Heo, S.J., Mitros, T., Munoz, D.P., Boffelli, D., et al. (2016). Selection-free genome editing of the sickle mutation in human adult hematopoietic stem/progenitor cells. Sci Transl Med 8, 360ra134.

Encode Project Consortium (2012). An integrated encyclopedia of DNA elements in the human genome. Nature 489, 57–74.

Gasperini, M., Tome, J.M., and Shendure, J. (2020). Towards a comprehensive catalogue of validated and target-linked human enhancers. Nat Rev Genet.

Gisler, S., Goncalves, J.P., Akhtar, W., de Jong, J., Pindyurin, A.V., Wessels, L.F.A., and van Lohuizen, M. (2019). Multiplexed Cas9 targeting reveals genomic location effects and gRNA-based staggered breaks influencing mutation efficiency. Nat Commun 10, 1598.

Goodarzi, A.A., Noon, A.T., Deckbar, D., Ziv, Y., Shiloh, Y., Lobrich, M., and Jeggo, P.A. (2008). ATM signaling facilitates repair of DNA double-strand breaks associated with heterochromatin. Mol Cell 31, 167–177.

Gottlieb, T.M., and Jackson, S.P. (1993). The DNA-dependent protein kinase: requirement for DNA ends and association with Ku antigen. Cell 72, 131–142.

Hendel, A., Kildebeck, E.J., Fine, E.J., Clark, J., Punjya, N., Sebastiano, V., Bao, G., and Porteus, M.H. (2014). Quantifying genome-editing outcomes at endogenous loci with SMRT sequencing. Cell Rep 7, 293–305.

Hustedt, N., and Durocher, D. (2016). The control of DNA repair by the cell cycle. Nat Cell Biol 19, 1–9.

Iliakis, G., Murmann, T., and Soni, A. (2015). Alternative end-joining repair pathways are the ultimate backup for abrogated classical non-homologous end-joining and homologous recombination repair: Implications for the formation of chromosome translocations. Mutat Res Genet Toxicol Environ Mutagen 793, 166–175.

Jakob, B., Splinter, J., Conrad, S., Voss, K.O., Zink, D., Durante, M., Lobrich, M., and Taucher-Scholz, G. (2011). DNA double-strand breaks in heterochromatin elicit fast repair protein recruitment, histone H2AX phosphorylation and relocation to euchromatin. Nucleic Acids Res 39, 6489–6499.

Janssen, A., Breuer, G.A., Brinkman, E.K., van der Meulen, A.I., Borden, S.V., van Steensel, B., Bindra, R.S., LaRocque, J.R., and Karpen, G.H. (2016). A single double-strand break system reveals repair dynamics and mechanisms in heterochromatin and euchromatin. Genes Dev 30, 1645–1657.

Jasin, M., and Haber, J.E. (2016). The democratization of gene editing: Insights from site-specific cleavage and double-strand break repair. DNA Repair (Amst) 44, 6–16.

Jeggo, P.A., and Downs, J.A. (2014). Roles of chromatin remodellers in DNA double strand break repair. Exp Cell Res 329, 69–77.

Jensen, K.T., Floe, L., Petersen, T.S., Huang, J., Xu, F., Bolund, L., Luo, Y., and Lin, L. (2017). Chromatin accessibility and guide sequence secondary structure affect CRISPR-Cas9 gene editing efficiency. FEBS Lett 591, 1892–1901.

Kallimasioti-Pazi, E.M., Thelakkad Chathoth, K., Taylor, G.C., Meynert, A., Ballinger, T., Kelder, M.J.E., Lalevee, S., Sanli, I., Feil, R., and Wood, A.J. (2018). Heterochromatin delays CRISPR-Cas9 mutagenesis but does not influence the outcome of mutagenic DNA repair. PLoS Biol 16, e2005595.

Kalousi, A., and Soutoglou, E. (2016). Nuclear compartmentalization of DNA repair. Curr Opin Genet Dev 37, 148–157.

Lee, Y.H., Kuo, C.Y., Stark, J.M., Shih, H.M., and Ann, D.K. (2013). HP1 promotes tumor suppressor BRCA1 functions during the DNA damage response. Nucleic Acids Res 41, 5784–5798.

Leemans, C., van der Zwalm, M.C.H., Brueckner, L., Comoglio, F., van Schaik, T., Pagie, L., van Arensbergen, J., and van Steensel, B. (2019). Promoter-Intrinsic and Local Chromatin Features Determine Gene Repression in LADs. Cell 177, 852–864 e814.

Lemaitre, C., Grabarz, A., Tsouroula, K., Andronov, L., Furst, A., Pankotai, T., Heyer, V., Rogier, M., Attwood, K.M., Kessler, P., et al. (2014). Nuclear position dictates DNA repair pathway choice. Genes Dev 28, 2450–2463.

Lin, S., Staahl, B.T., Alla, R.K., and Doudna, J.A. (2014). Enhanced homology-directed human genome engineering by controlled timing of CRISPR/Cas9 delivery. Elife 3, e04766.

Mateos-Gomez, P.A., Gong, F., Nair, N., Miller, K.M., Lazzerini-Denchi, E., and Sfeir, A. (2015). Mammalian polymerase theta promotes alternative NHEJ and suppresses recombination. Nature 518, 254–257.

McVey, M., and Lee, S.E. (2008). MMEJ repair of double-strand breaks (director’s cut): deleted sequences and alternative endings. Trends Genet 24, 529–538.

Mitrentsi, I., Yilmaz, D., and Soutoglou, E. (2020). How to maintain the genome in nuclear space. Curr Opin Cell Biol 64, 58–66.

Mladenov, E., Magin, S., Soni, A., and Iliakis, G. (2016). DNA double-strand-break repair in higher eukaryotes and its role in genomic instability and cancer: Cell cycle and proliferation-dependent regulation. Semin Cancer Biol 37-38, 51–64.

Montague, T.G., Cruz, J.M., Gagnon, J.A., Church, G.M., and Valen, E. (2014). CHOPCHOP: a CRISPR/Cas9 and TALEN web tool for genome editing. Nucleic Acids Res 42, W401–407.

Okamoto, S., Amaishi, Y., Maki, I., Enoki, T., and Mineno, J. (2019). Highly efficient genome editing for single-base substitutions using optimized ssODNs with Cas9-RNPs. Sci Rep 9, 4811.

Ott, C.J., Federation, A.J., Schwartz, L.S., Kasar, S., Klitgaard, J.L., Lenci, R., Li, Q., Lawlor, M., Fernandes, S.M., Souza, A., et al. (2018). Enhancer Architecture and Essential Core Regulatory Circuitry of Chronic Lymphocytic Leukemia. Cancer Cell 34, 982–995 e987.

Pfister, S.X., Ahrabi, S., Zalmas, L.P., Sarkar, S., Aymard, F., Bachrati, C.Z., Helleday, T., Legube, G., La Thangue, N.B., Porter, A.C., et al. (2014). SETD2-dependent histone H3K36 trimethylation is required for homologous recombination repair and genome stability. Cell Rep 7, 2006–2018.

Pokusaeva, V.O., Diez, A.R., Espinar, L., and Filion, G.J. (2019). Strand asymmetry influences mismatch repair during single-strand annealing. 847160.

Redwood, A.B., Perkins, S.M., Vanderwaal, R.P., Feng, Z., Biehl, K.J., Gonzalez-Suarez, I., Morgado-Palacin, L., Shi, W., Sage, J., Roti-Roti, J.L., et al. (2011). A dual role for A-type lamins in DNA double-strand break repair. Cell Cycle 10, 2549–2560.

Richardson, C.D., Kazane, K.R., Feng, S.J., Zelin, E., Bray, N.L., Schafer, A.J., Floor, S.N., and Corn, J.E. (2018). CRISPR-Cas9 genome editing in human cells occurs via the Fanconi anemia pathway. Nat Genet 50, 1132–1139.

Richardson, C.D., Ray, G.J., DeWitt, M.A., Curie, G.L., and Corn, J.E. (2016). Enhancing homology-directed genome editing by catalytically active and inactive CRISPR-Cas9 using asymmetric donor DNA. Nat Biotechnol 34, 339–344.

Riesenberg, S., Chintalapati, M., Macak, D., Kanis, P., Maricic, T., and Paabo, S. (2019). Simultaneous precise editing of multiple genes in human cells. Nucleic Acids Res 47, e116.

Ryu, T., Spatola, B., Delabaere, L., Bowlin, K., Hopp, H., Kunitake, R., Karpen, G.H., and Chiolo, I. (2015). Heterochromatic breaks move to the nuclear periphery to continue recombinational repair. Nat Cell Biol 17, 1401–1411.

Sartori, A.A., Lukas, C., Coates, J., Mistrik, M., Fu, S., Bartek, J., Baer, R., Lukas, J., and Jackson, S.P. (2007). Human CtIP promotes DNA end resection. Nature 450, 509–514.

Schmidl, C., Rendeiro, A.F., Sheffield, N.C., and Bock, C. (2015). ChIPmentation: fast, robust, low-input ChIP-seq for histones and transcription factors. Nat Methods 12, 963–965.

Schmidt, D., Wilson, M.D., Spyrou, C., Brown, G.D., Hadfield, J., and Odom, D.T. (2009). ChIP-seq: using high-throughput sequencing to discover protein-DNA interactions. Methods 48, 240–248.

Schwalb, B., Michel, M., Zacher, B., Fruhauf, K., Demel, C., Tresch, A., Gagneur, J., and Cramer, P. (2016). TT-seq maps the human transient transcriptome. Science 352, 1225–1228.

Scully, R., Panday, A., Elango, R., and Willis, N.A. (2019). DNA double-strand break repair-pathway choice in somatic mammalian cells. Nat Rev Mol Cell Biol 20, 698–714.

Shah, R.N., Grzybowski, A.T., Cornett, E.M., Johnstone, A.L., Dickson, B.M., Boone, B.A., Cheek, M.A., Cowles, M.W., Maryanski, D., Meiners, M.J., et al. (2018). Examining the Roles of H3K4 Methylation States with Systematically Characterized Antibodies. Mol Cell 72, 162–177 e167.

Shen, M.W., Arbab, M., Hsu, J.Y., Worstell, D., Culbertson, S.J., Krabbe, O., Cassa, C.A., Liu, D.R., Gifford, D.K., and Sherwood, R.I. (2018). Predictable and precise template-free CRISPR editing of pathogenic variants. Nature 563, 646–651.

Solovei, I., Wang, A.S., Thanisch, K., Schmidt, C.S., Krebs, S., Zwerger, M., Cohen, T.V., Devys, D., Foisner, R., Peichl, L., et al. (2013). LBR and lamin A/C sequentially tether peripheral heterochromatin and inversely regulate differentiation. Cell 152, 584–598.

Soria, G., and Almouzni, G. (2013). Differential contribution of HP1 proteins to DNA end resection and homology-directed repair. Cell Cycle 12, 422–429.

Sun, Y., Jiang, X., Xu, Y., Ayrapetov, M.K., Moreau, L.A., Whetstine, J.R., and Price, B.D. (2009). Histone H3 methylation links DNA damage detection to activation of the tumour suppressor Tip60. Nat Cell Biol 11, 1376–1382.

Tsouroula, K., Furst, A., Rogier, M., Heyer, V., Maglott-Roth, A., Ferrand, A., Reina-San-Martin, B., and Soutoglou, E. (2016). Temporal and Spatial Uncoupling of DNA Double Strand Break Repair Pathways within Mammalian Heterochromatin. Mol Cell 63, 293–305.

van Overbeek, M., Capurso, D., Carter, M.M., Thompson, M.S., Frias, E., Russ, C., Reece-Hoyes, J.S., Nye, C., Gradia, S., Vidal, B., et al. (2016). DNA Repair Profiling Reveals Nonrandom Outcomes at Cas9-Mediated Breaks. Mol Cell 63, 633–646.

van Schaik, T., Vos, M., Peric-Hupkes, D., and van Steensel, B. (2019). Cell cycle dynamics of lamina associated DNA. 2019.2012.2019.881979.

Villarreal, D.D., Lee, K., Deem, A., Shim, E.Y., Malkova, A., and Lee, S.E. (2012). Microhomology directs diverse DNA break repair pathways and chromosomal translocations. PLoS Genet 8, e1003026.

Vogel, M.J., Peric-Hupkes, D., and van Steensel, B. (2007). Detection of in vivo protein-DNA interactions using DamID in mammalian cells. Nat Protoc 2, 1467–1478.

Yeh, C.D., Richardson, C.D., and Corn, J.E. (2019). Advances in genome editing through control of DNA repair pathways. Nat Cell Biol 21, 1468–1478.

Zorita, E., Cusco, P., and Filion, G.J. (2015). Starcode: sequence clustering based on all-pairs search. Bioinformatics 31, 1913–1919.

